# Paternal CHH methylation potentiates stress responses against Pseudomonas syringae in Arabidopsis progenies

**DOI:** 10.1101/2023.05.10.540197

**Authors:** Yongping Li, Tianjia Liu, Even Yee Man Leung, Xusheng Zhao, Guopeng Zhu, Danny W-K Ng

## Abstract

Systemic acquired resistance (SAR) is an induced immune mechanism in plants, involving epigenetic regulation by chromatin remodeling and DNA methylation, which can be inherited to progeny following stress exposure. Intersexual epigenetic conflict sometimes leads to unequal expression of maternal and paternal alleles in offspring, resulting in parent-of-origin effects of inheritance. To better understand the parental contributions of epialleles in plant defense, isogenic *Arabidopsis* parental lines were mock-treated (M) and *Pseudomonas syringae* (*Pst*)-treated (P) for reciprocal crosses to produce F1 progenies (MP, PM). Together with their self-fertilized F1 descendants (MM, PP), the genome-wide inherited DNA methylation and transcriptomic changes against *Pst* were analyzed. F1 descendants shared widespread DNA methylation and transcriptional changes at transposable elements (TEs) and genes. The confrontation of epigenomes triggers the reprogramming of DNA methylation in reciprocal crosses, resulting in transgressive segregation that also shows the parental effect of *Pst* treatment. Compared to PM, the MP (*Pst*-primed paternal genome) was found to contributes to CHH hypermethylation, which is associated with processes in plant-pathogen interaction, including carbohydrate metabolism, glutathione metabolism and stronger translation process, which potentially contribute to improved disease resistance in MP in response to *Pst* challenge. Our data suggested a parent-of-origin effect of defense priming that contributes differently toward improved defense response in progenies.

## Introduction

Being sessile, plants have evolved various signaling mechanisms to cope with the rapidly changing environments and biotic stresses throughout their life cycle (Bilichak and Kovalchuk, 2013; Jones and Dangl, 2006; Pieterse et al., 2009). One such mechanism is systemic acquired resistance (SAR) that enables the whole plant to mount defense against a pathogen following an initial localized exposure to the pathogen (Fu and Dong, 2013). Over the last decade, defense priming for enhanced responses to abiotic or biotic stress within a single generation has been well studied (Borges and Sandalio, 2015; Conrath et al., 2001). Accumulating evidence have suggested that SAR not only could prime plants to develop a more rapid response and resistance against subsequent pathogen challenges in the parental generation, such primed state was indeed “recorded” and be able to pass onto the next generation, thereby enhancing their progenies’ response to stresses and survival (Chinnusamy and Zhu, 2009; Conrath et al., 2002). There is always a trade-off between growth and defense during plant development (Todesco et al., 2010). Constitutive expression of extra copies of the disease resistance (R) gene, *Resistance to Pseudomonas syringae protein 3* (*RPM1*), in *Arabidopsis* has led to lower shoot biomass and seed production (Tian et al., 2003). Therefore, defense priming poses an evolutionary advantage in maximizing resource allocations for growth and development while allowing the plant to elicit an enhanced defense response when needed (Lozano-Duran et al., 2013; Neilson et al., 2013; Paul-Victor et al., 2010).

Parental effect, especially through maternal provisioning, has been shown to contribute to seed sizes and germination in the progenies (Costa et al., 2012). While epigenetic contribution to transgenerational inheritance is evident, it is unclear if parental effects play a role in governing such pattern of inheritance in defense response. In crop breeding, parental selection is one of the major decisions that plant breeders have to make in order to maximize the production and quality of recombinant offspring (in another word, heterosis). When genomes from two species are brought together through hybridization, genome-wide changes such as histone modifications, DNA methylation and transposons reactivation are expected to contribute to the development of heterotic phenotypes (Chen, 2010; Doyle et al., 2008). Maternally- and paternally-controlled traits were found to associate with different mode of inheritance of gene expression in rapeseed hybrids. For maternally-controlled traits, their expressions are additive in the hybrid while a dominant inheritance was found for traits that are controlled by the paternal parents (Xing et al., 2014). Moreover, it has been found that in carrot, hybrid characteristics were influenced by paternal effect rather than by maternal effect (Grebenstein et al., 2013) and such parent-of-origin effect could be attributed to epigenetic controls (Curley et al., 2011). Recently, it has been shown that paternal transgenerational immune priming can confer immune protection in offspring of red flour beetle (Eggert et al., 2014). Therefore, the selection of maternal and paternal parents in breeding potentially could affect hybrid performance in breeding.

DNA methylation is an epigenetic modification that is widespread in plants and can be separated into three sequence context (CG, CHG, and CHH, where H denotes A, T or C), mainly to silence repeats and transposable elements (TEs) in heterochromatic regions (Law and Jacobsen, 2010; Zhang et al., 2018). In *Arabidopsis*, methylation is maintained by METHYLTRANSFERASE1 (MET1), CHROMOMETHYLASE3 (CMT3), and CMT2 for CG, CHG and CHH sites, respectively (Bartee et al., 2001; Finnegan et al., 1996; Finnegan and Dennis, 1993; Kato et al., 2003; Lindroth et al., 2001). The CMT2 mediates CHH methylation of transposon element (TE) in pericentromeric regions (Gouil and Baulcombe, 2016; Stroud et al., 2014). The RNA-directed DNA Methylation (RdDM) pathway is responsible for *de novo* methylation, a process that is most clearly observed at CHH sites. Besides, RdDM can help the host to respond to biotic or abiotic challenges, and also affect germ cell specification and parent-specific gene expression (Barber et al., 2012; Matzke and Mosher, 2014). DNA methylation is also involved in priming; the transposable elements and their surrounding sequences showed CHH hypomethylation in *rdd* (*ros1/dml2/dml3*) mutants, resulting in susceptibility of plants to pathogens (Zhou et al., 2014).

In this study, we performed a comprehensive genome-wide investigation of the epigenome and transcriptome of self-fertilized descendants of the mock-treated (MM), *Pst*-treated parents (PP) and their reciprocally crossed F1 progenies (MP and PM). Differential response to *Pst* challenge was observed among the F1 descendants. Widespread DNA methylation and transcriptional changes at transposable elements (TEs) and genes were observed among the four descendants. Here, we aimed to evaluate the effects of parental *Pst* treatment on intergenerational TE and gene expression changes among F1 progenies showing differential responses the pathogen. Primed genome induces rapid changes in chromatin organization, leading to a more rapid disease resistance response against *Pst* infection. Primed PP alters the dynamic of defense-responsive genes, contributing to its improved resistance against *Pst*. For reciprocal crosses (MP and PM), epigenome confrontation triggers reprogramming of DNA methylation, leading to transgressive segregation and also shows the parental effect of *Pst* treatment. It is worth noting that the *Pst*-primed paternal genome contributes to CHH hypermethylation, which potentially contribute to improved disease resistance in MP. These findings will facilitate the selection of maternal and paternal epigenetic variation for breeding.

## Results

### Improved defense response in F1 descendants of *Pst*-primed parents

To understand if parent-of-origin effect has contribution in conferring transgenerational defense priming in *Arabidopsis*, we have created F1 descendants from isogenic *Arabidopsis thaliana* (L*er* ecotype) through reciprocal crosses between a mock-treated and a pathogen-treated parents (Fig. 1A). In addition, seeds from the selfed-parents were collected as controls. Among the F1 descendants, a difference in disease symptoms was observed between self-fertilized descendants (MM and PP) of the mock-treated (M) and the *Pst-*treated (P) parents. Moreover, a more pronounced chlorotic phenotype was observed in MM than that in PP after 2 days post infection (dpi) (Fig. 1B-C); and this is consistent with the findings previously reported by Luna et al. (2012) (Luna et al., 2012). In the reciprocal descendants from cross-fertilization between the M and P parents, MP and PM plants also displayed less severe disease symptoms when compared to MM (Fig. 1B). The degree of disease symptoms of the F1 descendants was consistent with the endophytic bacterial growth in the leaf tissues (Fig. 1C). Therefore, these data confirmed a priming effect for enhanced stress responses in the progenies when the parental plants were exposed to stresses. Interestingly, between MP and PM, MP plants showed more resistance to *Pst* when compared to PM (Fig. 1C). Such transgenerational inheritance reflects a potential parent-of-origin effect to which the paternally-primed parent conferred a higher disease resistance in the descendants than that of the maternally-primed parent. Therefore, these data supported that uni-parental stress-priming is sufficient to confer an improved defense response in the progenies.

**Fig. 1.**
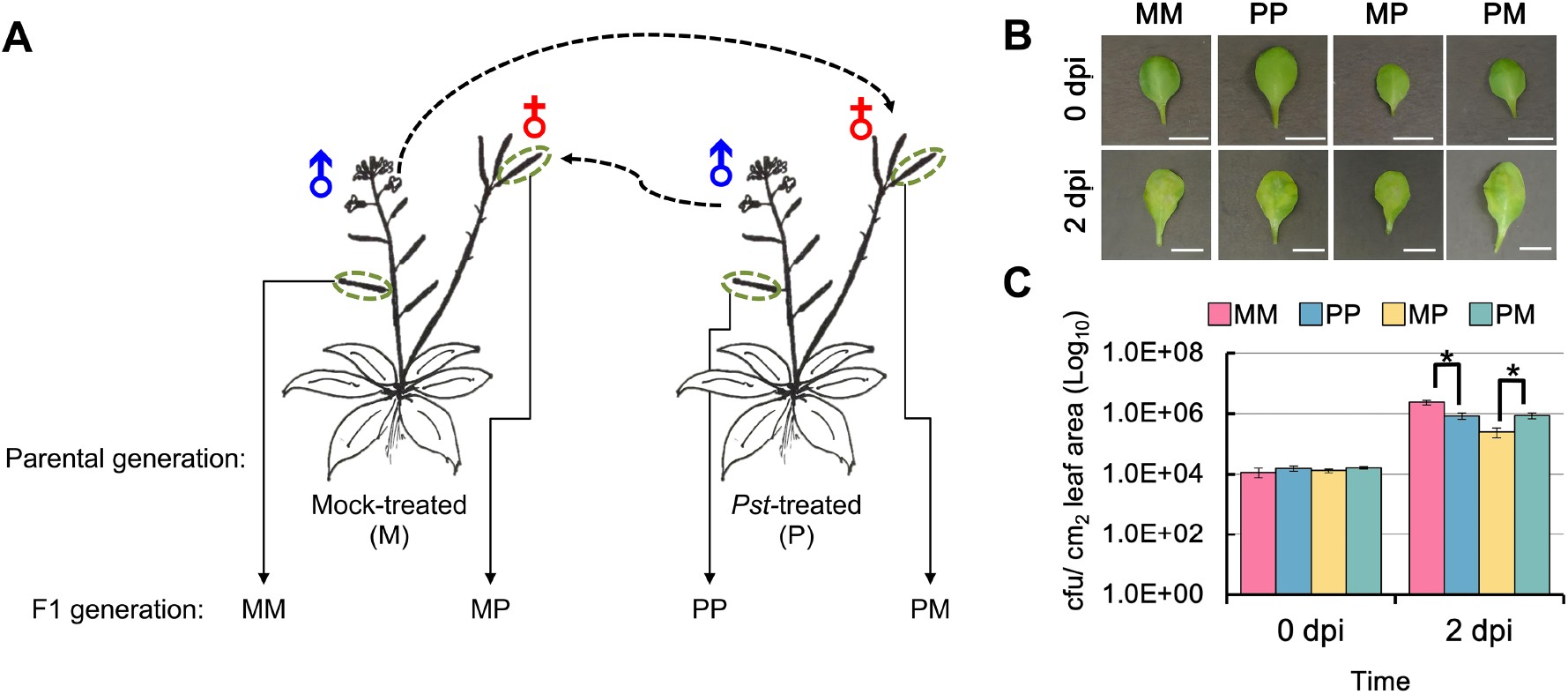
Parental and parent-of-origin immune priming for defense responses against *Pseudomonas syringae* in *Arabidopsis.* (A) Schematic design of crossing experiments. Parental plants were subjected to either mock-treatment (M) or inoculation-treatment with *Pseudomonas syringae* pv. *tomato* DC3000 (*Pst*DC3000) (P) during their vegetative growth prior bolting. Descendants from the corresponding selfed parents (MM and PP), and their reciprocally crossed progenies (MP and PM) were collected for subsequent analyses. (B) Mature leaves from the descendant plants were treated with *Pst*DC3000 (5×10^6^ CFU/mL, OD_600_ = 0.01) by syringe infiltration. Representative leaf showing disease symptoms at 2 days post infection (dpi) were captured. Scale bar = 1 cm. (C) Leaf disks were collected at indicated time point and the bacterial titers (CFU/ cm^2^ leaf tissue) were determined. Data are mean ± SE from three replicates. Asterisks represent significant difference at P < 0.05 in t-test.

### Paternal primed genome triggers CHH hypermethylation in TE regions

To investigate the impacts of parental *Pst* treatment on DNA methylation states in the F1 descendants, we generated single-base resolution maps of DNA methylation for the four F1 descendants, MM, PP, MP, and PM (Fig. 1A; Supplemental Table S1). For each library, at least 50M pair-end read (read length = 100bp) were produced, approximately 54-60% of the reads were mapped to the *Arabidopsis* genome using Bismark (Krueger and Andrews, 2011). In the F1 descendants, the total number of methylated cytosines was 24, 8, and 3% in the CG, CHG and CHH contexts, respectively (Supplemental Table S1). First, the genome-wide methylation levels (the genome was divided into 100bp bins) were calculated for each sequenced descendant. The self-fertilized descendant of the mock-treated (MM) was used as the baseline to assess changes to other descendants, which revealed median methylation level of 86.0, 45.0, and 16.0% for CG, CHG, and CHH, respectively. Increased CHH methylation in MP (18.0%) relative to other descendants was readily apparent (Fig. S1), and CHG methylation was decreased slightly with the order of MP, PM and PP.

We observed similar changes when characterizing the global methylation of protein-coding genes and TEs region in CG, CHG, and CHH cytosine context for these descendants. All four descendants showed a similar pattern of CG and CHG methylation in both protein-coding genes and TEs (Fig. 2A). In contrast, CHH methylation level at gene regions showed a slight increase trend with the order of MM, MP, PM and PP, suggesting an effect of DNA methylation of *Pst* treatment in the parental genome. Consistent with results obtained from 100 bp windows at whole-genome methylation level (Fig. S1), CHH methylation increased significantly at the TE and its surroundings regions in MP (Fig. 2A). The average DNA methylation levels of TEs increases with increasing distances from the nearby genes (Fig. S2). The *Arabidopsis* genome contains repetitive sequences and has an accumulation of transposable elements in pericentromeric regions of the chromosomes (Simon et al., 2015). These regions have a higher density of TEs compared to euchromatin. Therefore, to determine the genome-wide impact of hypermethylation observed in the F1 descendants, we calculated the DNA methylation level of the CG, CHG and CHH contexts in all descendants (Fig. S2). The results showed that the genome-wide CHH hypermethylation occurred throughout the chromosome of MP, with the greatest increases in TE-enriched regions. Collectively, these results indicate that increases in CHH methylation in MP occur across the entire genome and correlate with the abundance of TE. However, both CG and CHG methylation are maintained in a symmetrical manner with high fidelity, suggesting that methylation in CG and CHG contexts are more stable than that in CHH context.

**Fig. 2.**
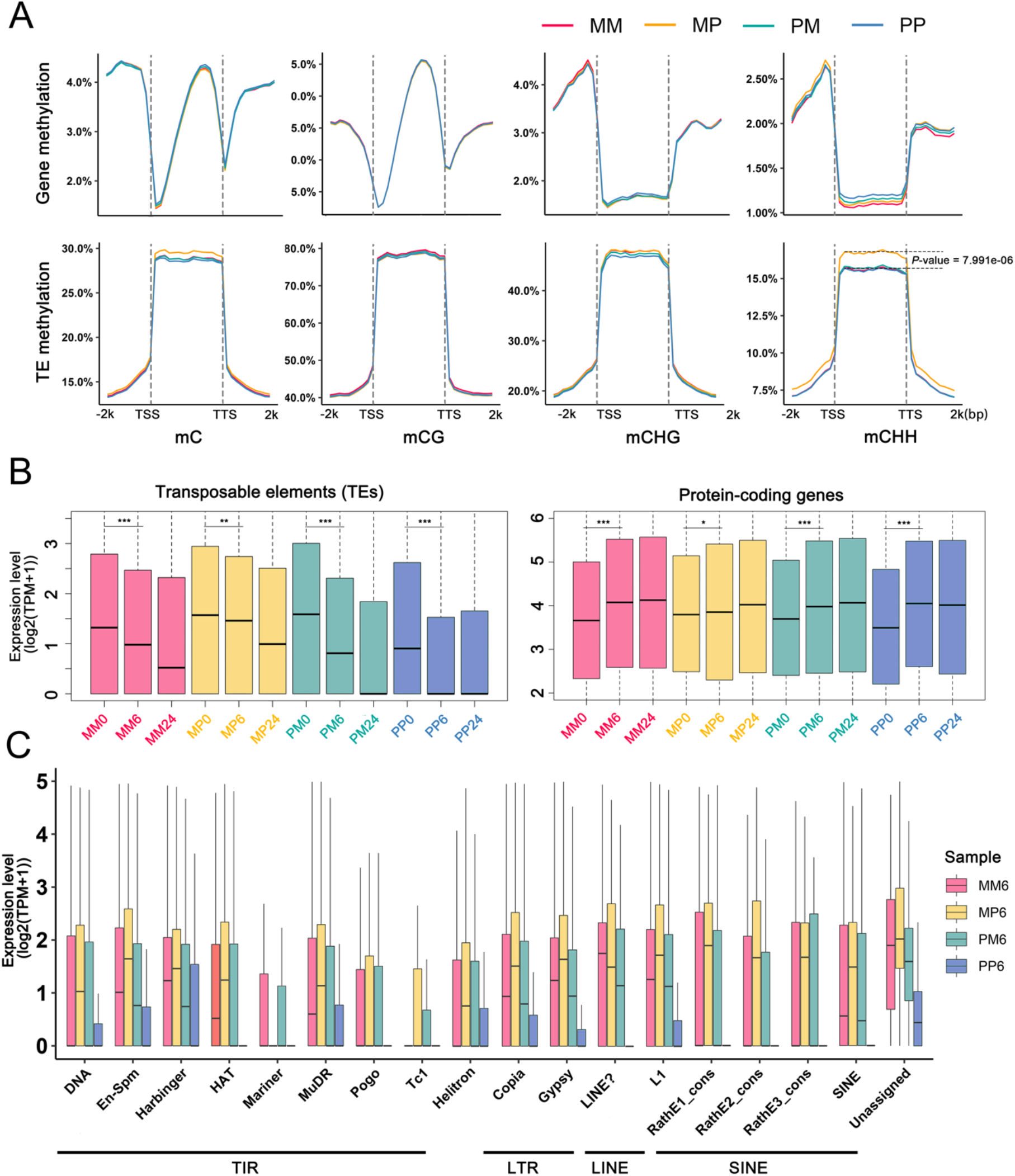
Methylation patterns and gene/TE expression in F1 descendants. (A) DNA methylation profiles of mCG, mCHG, mCHH and mC surrounding genes (upper panel) and TEs (lower panel) in MM, MP, PM and PP. Transcription start site (TSS) and transcription termination site (TTS) are indicated. (*P*-value < 0.001, as determined using the t-test). (B) Read density along with total reads from four descendant RNA-Seq libraries. Expressed genes (17,583) and TEs (12,468) with TPM ≥ 2 in more than one sample were used for box plots. The top, middle, and bottom lines of the box indicate the 25th, 50th, and 75th percentiles, respectively. (*: *P* < 0.05 (significant), **: *P* < 0.01 (highly significant); ***: *P* < 0.001 (extremely significant), as determined using the t-test). (C) Expression level of TEs among different TE superfamilies in MM, MP, PM and PP at 6h after *Pst* treatment.

### *Pst* treatment silences TEs

Parent-of-origin effects are often associated with maternal and paternal inheritance of DNA methylation patterns (Ferguson-Smith, 2011; Raissig et al., 2011), and methylation variation of transposable elements (TEs) affects gene expression levels (Zhang et al., 2015). It has been shown that epigenetic states of TEs was dynamically altered in response to biotic stress (Dowen et al., 2012). Here, we analyzed expression landscapes of TEs and protein coding genes across all F1 descendants using transcriptome data, quantifying the expression abundance of TE and gene transcripts across various F1 genomes at 0, 6 and 24 hours in response to *Pst* challenge (Fig. 2B). Using mRNA-seq, we were able to recover expressed transcripts of 12,468 and 17,583 (TPM ≥ 2 in more than one sample) out of the 31,189 and 27,655 TEs and protein-coding genes annotated in Araport11, respectively. Before *Pst* treatment, global expression levels of TEs and protein coding genes are higher in reciprocal lines (MP and PM) than that in the selfed-lines (MM and PP). After *Pst* treatment, global expression of TEs dramatically declined in PP, reaching to a very low level at 6 hours. PM reached a similar low level of TEs expression at 24 hours after *Pst* infection. Although expression level of TEs in MM and MP also showed a declining trend, TEs expression in both lines was maintained at a relatively high level at 24 hours after *Pst* infection, with the highest level being detected in MP (Fig. 2B). Contrary to TEs expression, protein-coding gene showed an opposite trend of expression after *Pst* treatment. Overall, gene expression was upregulated with the progression of *Pst* infection. Among the F1 descendants, gene expression displayed a greater variation in MM, PM and PP before/after *Pst*, suggesting a more dynamic response to transcriptome changes in these descendants. In contrast, the expression levels of both TEs and protein-coding genes in MP were demonstrated to be more stable. According to the result above, the increased CHH methylation could be associated with transcriptional activation program by maintaining a higher expression level of TEs before/after *Pst* infection.

It is interesting to note that TEs expression was maintained at a higher level in MP when compared to PM, MM and PP at 6 hours after *Pst* treatment. To further understand the effect of *Pst* treatment on TEs expression, we calculated the global expression level of different TE superfamilies at 6 hours after *Pst* treatment (Fig. 2C). The response of TEs in F1 descendants was highly specific among different TE superfamilies and their expressions were sorted into different TE classes according to recent study (Quesneville, 2020). First, the Class II with order of TIR (including the DNA, EN-Spm, Harbinger, hAT, Mariner, MuDR, Pogo and Tc1 superfamilies), the expression of most superfamilies within this class is consistent with the global expression. But we found that there are two TE superfamilies showing completely different expression patterns with other superfamilies. For example, the Mariner superfamily was only highly expressed in MM and PM, suggesting a paternal effect on the methylation of the transposon elements. Besides, the Tc1 superfamily was found to be highly expressed only in the two reciprocal lines (MP and PM). Interestingly, these two TE superfamilies share a common amino acid motif called the “DDE/D” signature. In protozoan, the Tc1/Mariner transposable element family shapes genetic variation and gene expression in the *Trichomonas vaginalis* (Bradic et al., 2014). Although this family of TE has been identified in plants, their functions are less studied (Liu and Yang, 2014). Other TE superfamilies show similar expression patterns to global TE expression.

### Parental *Pst* treatment leads to transgenerational DMRs among F1 descendants

To assess the transgenerational effect of parental *Pst* treatment on DNA methylation among the F1 descendants, MM was used as reference to identify differential methylated regions (DMRs) among PP, MP and PM under normal growth condition (Fig. 3A; Table S3). Among the three ‘*Pst*-primed’ F1, PP contained the largest number of hypermethylated (hyperDMRs) in CG (776) and CHG (249) contexts (Fig. 3A). Similarly, a greater number of hypomethylated DMRs (hypoDMRs) were identified in CG and CHG contexts (744 and 101, respectively). In contrast, DMRs in CHH contexts showed a higher abundance in both MP and PM than that in PP. Between MP and PM, MP showed a slightly higher number of CHH methylation, with 92 hyperDMRs and 71 hypoDMRs, respectively. Next, we examined genomic features overlapping with DMRs (Fig. 3B). We found that CG methylation predominantly overlaps with coding sequence (CDS) (72% for the hyperDMRs, 78% for the hypoDMRs; Fig. 3B). The distribution of hyperDMRs and hypoDMRs in the CHG context were similar in CDS, TE and intergenic regions (38%, 28% and 32% for hyperDMRs; 36%, 30% and 32% for hypoDMRs, respectively). In contrast, the CHH context mostly overlapped with the intergenic regions (46% and 62% for hyper- and hypoDMRs, respectively), next are overlapping with the TE (46% and 36% for hyper- and hypoDMRs, respectively), and to a lesser extent with the CDS. These results suggest that CG methylation potentially regulates gene expression, whereas CHH methylation regulates TE expression. Our finding is consistent with a previous study reporting that the CHH methylation at TE can undergo dynamic change during seed development and germination (Kawakatsu et al., 2017). In terms of the overall degree of methylation among the F1 descendants, hyperDMRs showed an increasing trend with the order MM < MP and PM < PP while hypoDMRs showed an opposite trend of methylation in all three contexts (CG, CHG, CHH) (Fig. 3C). Therefore, our methylation data show that parental *Pst* treatment has differential effects on descendants DNA methylation.

**Fig. 3.**
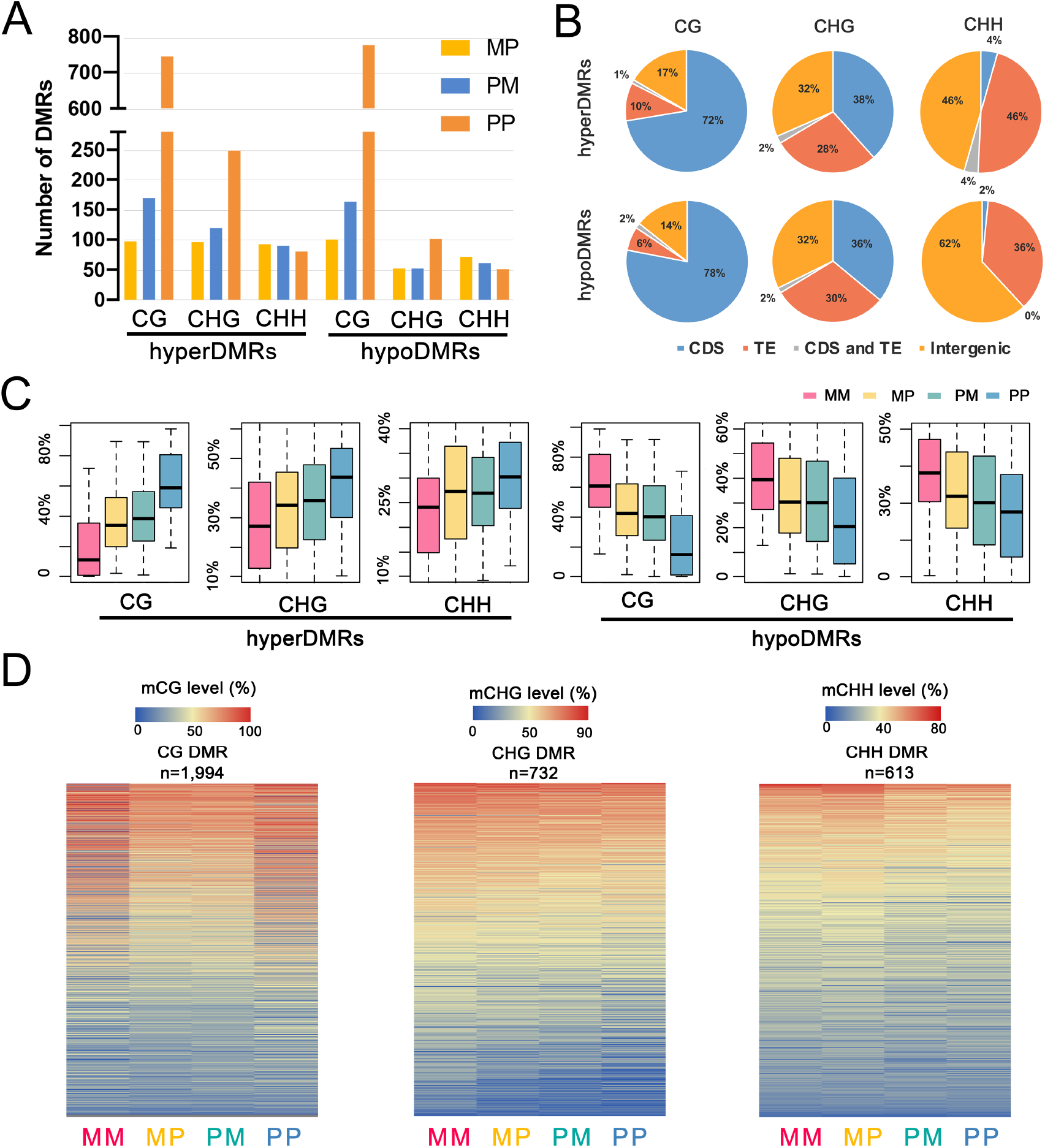
DMRs in F1 descendants. (A) Total number of DMRs (compared with MM) found in the three methylation contexts (CG, CHG, and CHH). Hyper- and hypoDMRs are shown. (B) The distribution of hyper- and hypoDMRs between MM and PP in genome elements. (C) DNA methylation level (%) at hyper- and hypoDMRs in CG, CHG and CHH contexts. between MM and PP. Regions were considered to be differentially methylated when the absolute differences of methylation were at least 0.4, 0.2, 0.2 for CG, CHG and CHH, respectively. (D) Heatmaps of methylation levels at CG DMRs, CHG DMRs, and CHH DMRs in F1 descendants. The total number of DMRs is labeled above each heatmap. The DMRs were sorted by methylation level.

To ensure the observed difference in DNA methylation among the F1 descendants is linked to parental *Pst* treatment, an unbiased assessment of DNA methylation changes was conducted. MethylC-seq reads were pooled together for an undirected identification of DMRs across all samples in the CG, CHG and CHH contexts (Fig. S4). In total, 2,884 CG, 1,108 CHG, and 893 CHH DMRs were identified throughout the entire genome (Fig. S4). After merging overlap DMRs from each comparison, a total of 1,994 CG, 732 CHG and 613 CHH DMRs were obtained (Fig. 3D). From the methylation level of DMRs, we found that the CG methylation level of MP and PM is the average of MM and PP, which is not observed in CHG and CHH context. CHH hypermethylation in MP was also observed in the level of CHH DMRs. Thus, the inheritance of CG methylation is very stable, whereas the level of CHH methylation is altered in the F1. Among the CG, CHG, and CHH DMRs occurring in intergenic regions, 65.4, 81.9, and 73.7% of them overlapped with TEs, respectively. In addition, 7.5% of CG DMRs, 14.1% of CHG DMRs, and 24.0% of CHH DMRs are located in promoter regions (Fig. S4). Among the CG, CHG, and CHH DMRs occurring in promoter regions, 21.0, 33.5, and 48.8% of them overlapped with TEs, respectively. 84.0, 43.1 and 24.0% of CG, CHG, and CHH DMRs are located in exon regions (Fig. S4). A larger number of CG DMRs were located in the gene region, in contrast, the CHH DMRs were mainly located in intergenic regions.

### Priming induces differential DNA methylation and gene expression changes in response to *Pst* in F1

Our high-resolution methylome uncovered a subset of genes or regions that gain or lose DNA methylation among F1 descendants when primed with *Pst* challenge in the parental generation (Fig. S5). In addition, increased expression of TEs was detected in the F1 descendants when compared to the unprimed line (MM). For instance, a transposable element gene named sadhu non-coding retrotransposon 3-2 (SADHU3-2), was completely demethylated in PP and reduced methylation was also detected in MP and PM when compared to MM (Fig. 4A). At transcriptional level, a high basal expression of SADHU3-2 was evident in PP, MP and PM and such high level of expression was maintained when the plants were challenged with *Pst* at 6 and 24 hours after infection (Fig. 4B). In some natural accessions, the SADHU3-2 allele is methylated and silenced, and is also involved a stable, meiotically transmissible epigenetic allele (Rangwala et al., 2006).

**Fig. 4.**
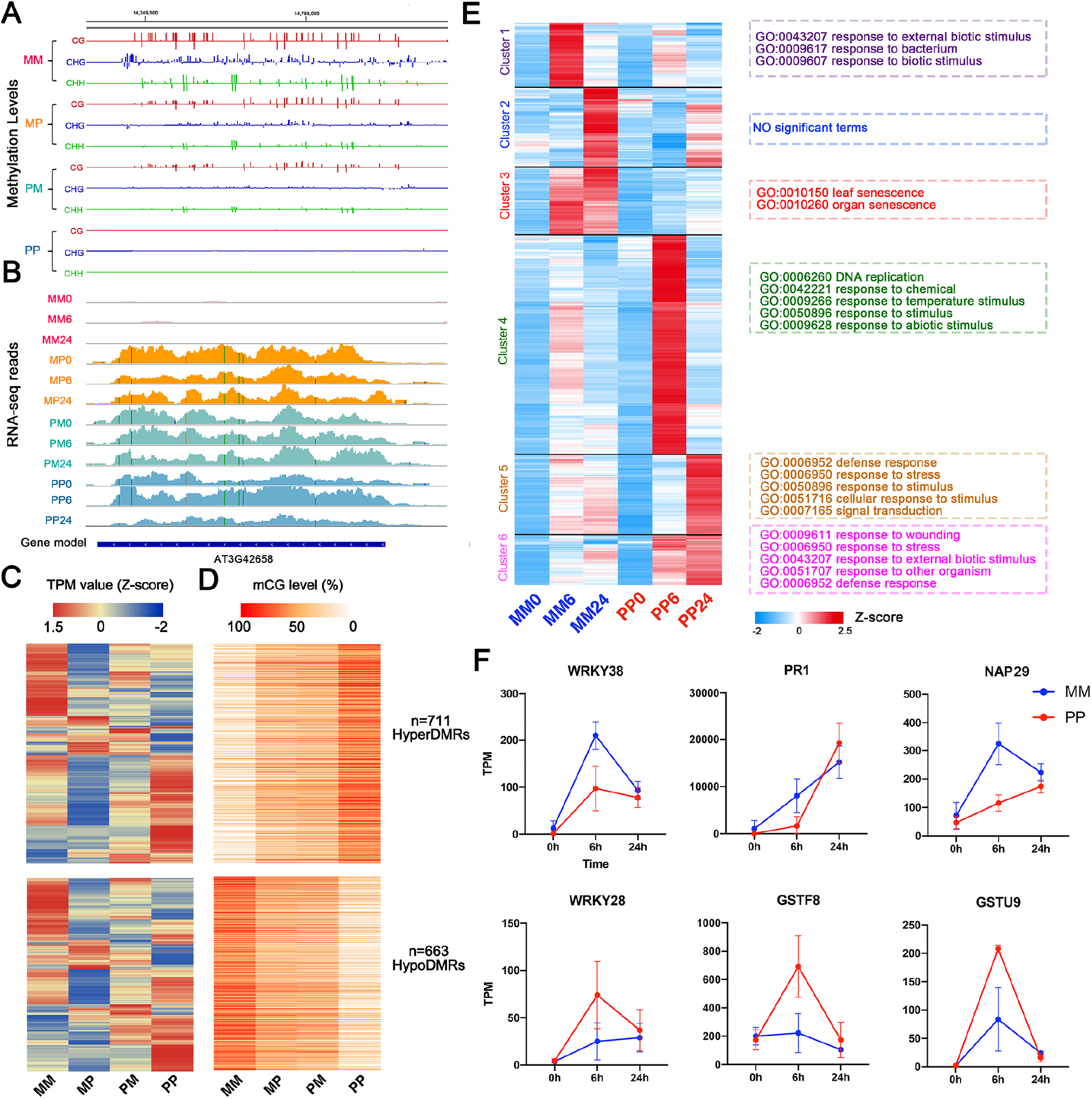
Priming effect induces changes in DNA methylation and gene expression for enhanced resistance against *Pst* infection. (A) Genome browser capturing the DNA methylation levels of sadhu3-2 in MM, MP, PM and PP. Gene models are represented in blue. CG methylation is shown in red, CHG in blue, and CHH in green. (B) IGV view of the SADHU3-2 RNA-Seq reads across the MM, MP, PM and PP at three time points. (C) Heatmaps showing the expression patterns of genes associated with CpG DMRs. Z-score obtained from averaged TPM of three biological replicates was used. (D) Heatmaps showing the CpG methylation level in corresponding overlapping CpG DMRs. The number of DMRs are labeled on the right side. (E) Hierarchical clustering of genes that are differentially expressed and GO enrichment results. (F) Examples of expression profiles of tissue-specific genes in our study. y-axis, transcript abundance; x-axis, time (h) after inoculation with *Pst*DC3000; error bars indicate SE.

To quantify gene expression changes associated with the changes in DNA methylation levels, two categories of genes were defined based on the correlations. In total, we identified 767 differentially expressed genes with 711 hyperDMRs (gaining at least 40% CpG methylation). In contrast, 703 genes were found to overlap with 663 hypoDMRs (losing at least 40% CpG methylation) (Fig. 4C-D). From the gene expression profiles, we found that most of the genes were highly expressed in MM or PP due to the transcription activation or repression, suggest that the expression profiles of genes are associated with hyper- or hypoDMRs (Fig. 4C). Genes that gain/loss DNA methylation will affect their transcriptional activity (Fig. 4C), suggesting a potential priming effect from *Pst* treatment in the parental generation. In order to determine the genome-wide transcriptome changes in F1 when challenged with *Pst*, DEGs between the untreated (0 hours) and *Pst*-treated (6 and 24 hours) plant samples in MM and PP were identified (Fig. S6; Table S4-S5). The number of up-regulated genes in PP was significantly more than that in MM, suggesting a priming effect for altered defense response and gene expression in F1 when the parental generations were exposed to *Pst.* PP showed improved resistance again *Pst* infection when compared to MM (Fig. 1C), genes highly expressed in MM6 (Fig. 4E) may involve in negative regulation of defense response. Indeed, it has been reported that *WRKY38* (Fig. 4F) is induced by *Pst* and overexpression of *WRKY38* leads to reduced disease resistance in *Arabidopsis* (Kim et al., 2008). Interestingly, we found genes that are highly expressed in both MM6 and MM24 (Cluster 3) were enriched in leaf and organ senescence GO terms. In *Arabidopsis*, inducible overexpression of AtNAP (Arabidopsis NAC domain containing protein) was found to cause precocious senescence (Gonzalez-Bayon et al., 2019; Guo and Gan, 2006). In our data, *AtNAP29* was found to be less induced in PP when compared to MM at 6 and 24 hours upon *Pst* challenge, reflecting an accelerated senescence pathway in MM upon infection by *Pst*.

Genes in cluster 4 were highly expressed in PP6 but not in MM6. Genes within this cluster may be involved in positive regulation of defense response and termed as primed genes. These genes were also highly expressed across the F1 descendants (Fig. S7). These genes are involved in DNA replication and response to chemical. DNA replication has been reported to be involved in histone modification, which maintains polycomb gene silencing in plants and the inheritance of the silencing memory from mother to daughter cells (Jiang and Berger, 2017). Overexpression of *WRKY28* activate jasmonic acid/ethylene pathway to defend *Arabidopsis* against oxalic acid and *Sclerotinia sclerotium* (Chen et al., 2013). WRKY28 also confers resistance to abiotic stress in *Arabidopsis* (Babitha et al., 2013). In PP, a more rapid and high induction of *AtWRKY28* was observed when compared to MM at 6 hours after *Pst* infection (Fig. 4F), reflecting an improved resistance against *Pst* in PP. Two glutathione S-transferase genes (*GSTF8* and *GSTU9*) are also included in cluster 4 (Fig. 4F), which are involved in toxic substances and oxidative stress (Lou et al., 2020). Genes highly expressed in PP24 (Cluster 5) were enriched for defense response, response to stress, response to stimulus and signal transduction. One of cluster 5 genes, *PR1*, a salicylic acid inducible maker for the SAR response, a delay but higher induction of *PR1* expression was detected in PP. Therefore, the altered dynamic of these defense-responsive genes in the primed PP could contribute to its improved resistance against *Pst*.

### Parental effect of gene expression changes in the F1 descendants

To further understand the possible parent-of-origin effects on disease resistance, we compared the *Pst-*induced transcriptomes of MP and PM against that in MM at 0, 6 and 24 hours after *Pst* challenge (Fig. 5A; Fig. S10; Table S7). Overall, we detected 348 (157 up- and 191 down-regulated) and 242 (148 up- and 94 down-regulated) genes to which their expression was altered at 6 hours and 24 hours respectively upon *Pst* treatment in MP. Since this set of genes resulted from the paternal *Pst-*priming (MP), we thus referred them as paternally primed genes (PPGs). In contrast, we have detected 32 (26 up- and 6 down-regulated) and 328 (244 up- and 84 down-regulated) genes with altered expression at 6 hours and 24 hours respectively in the reciprocally primed F1, PM (maternal *Pst-*primed). This set of genes thus referred as maternally primed genes (MPGs).

**Fig. 5.**
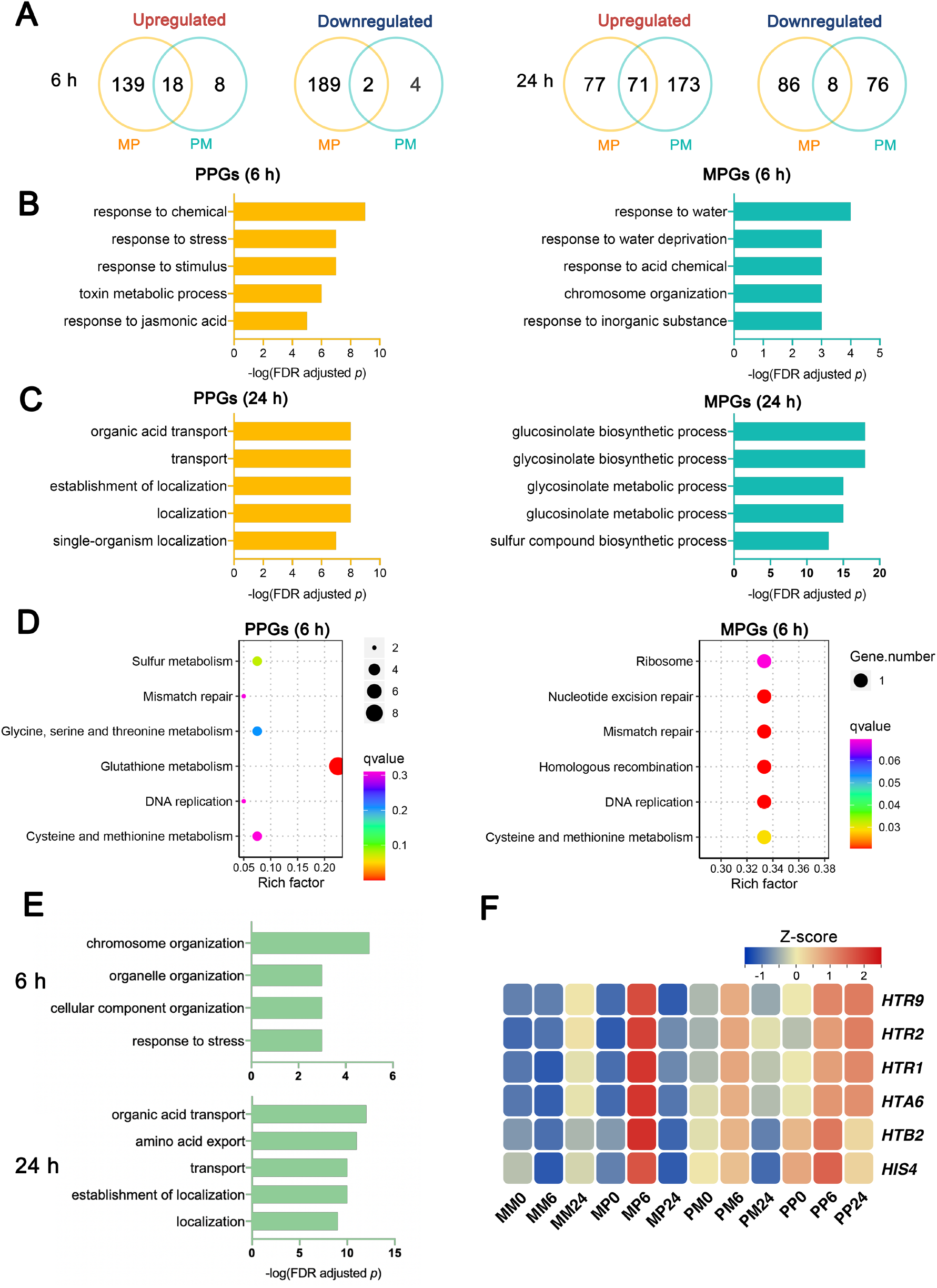
GO and KEGG analysis of PPGs and MPGs against *Pst* infection. (A)Venn diagrams showing the number of up- and down-regulated DEGs response to *Pst*DC3000 in MP and PM at 6 and 24 hours, when compared with MM, respectively. (B) GO-term enrichment for PPGs and MPGs identified at 6 hours, respectively. (C) GO-term enrichment for PPGs and MPGs identified 24 hours, respectively. (D) KEGG analysis of PPGs and MPGs at 6 hours. The statistical analysis was performed using a hypergeometric test. PPGs, paternal primed genes; MPGs, maternal primed genes. (E) GO-term enrichment for genes co-upregulated in MP and PM at 6 and 24 hours. (F) Heatmaps showing the RNA expression pattern of six histone-related genes.

To investigate the roles of these DEGs in the process of disease resistance, we performed GO enrichment analysis for each PPGs and MPGs in the reciprocally *Pst*-primed F1 descendants (Table S8). Results showed that different molecular pathways were enriched in MP and PM (Fig. 5B-C), PPGs and MPGs show different GO functional differences at different time points after *Pst* treatment. The PPGs in MP were enriched in biological GO categories that are involved in enhancing disease and stress responses, especially at 6 hours with biological processes including response to chemical, response to stress, response to stimulus and response to jasmonic acid. The MPGs in PM were enriched mainly in biological processes related to response to water, response to water deprivation, response to acid chemical, and chromosome organization at 6 hours. Notably, several glucosinolate and glycosinolate related GO terms were highly enriched among the MPGs at 24 hours, including glucosinolate biosynthetic process, glycosinolate biosynthetic process, glycosinolate metabolic process and glucosinolate. The result indicates that glucosinolate/ glycosinolate metabolism genes may be an important factor contributing to the disease resistance in PM.

We then used the Kyoto Encyclopedia of Genes and Genomes (KEGG) database to investigate potential PPG and MPG pathways. PPGs and MPGs at 6 hours were classified into known KEGG pathways. Consistent with the GO analysis results, PPGs and MPGs were also grouped into distinct KEGG pathways (Fig. 5D). Pathways of glutathione metabolism, cysteine and methionine metabolism, and DNA replication were enriched among the PPGs at 6 hours after *Pst* treatment. Among the MPGs, pathways of ribosome, nucleotide excision repair and mismatch repair were enriched.

### Primed genome shows enhanced expression of genes involved in chromatin organization rearrangement

Interestingly, we found that genes upregulated in both MP and PM were also enriched in distant biological processes at different time points. Genes upregulated at 6 hours showed enriched for GO terms related to chromosome organization, organelle organization, and cellular component (Fig. 5E). Besides, the core histone (CH) gene family (Fig. S12) is also enriched in 6 hours. Many histone-related genes were identified among the upregulated genes, such as *histone H2A 6* (*HTA6*), *histone B2* (*HTB2*) *histone H4* (*HIS4*) and three *histone 3.1* genes (*HTR1, HTR2 and HTR9*) (Fig. 5F). Previous study reported that heat stress can cause global 3D chromatin organization rearrangement with activation of TEs around rearranged regions in *Arabidopsis* (Sun et al., 2020). Furthermore, a recent study reported that H3K27me3 led to the interactions within polycomb-associated repressive domains, which induces a global reconfiguration of chromatin architecture (Huang et al., 2021). In this study, the polycomb gene related GO term (DNA replication) was found to be enriched in descendants which contain one or more primed genome (Fig. S8), suggesting a possible chromatin organization rearrangement in the primed genome. In contrast to this, several GO terms related to transport or localization were enriched at 24 hours, including organic acid transport, amino acid export, transport, and establishment of localization, these biological processes may involve in the later stage of *Pst* infection.

### CHH hypermethylation in MP improves disease resistance

In the reciprocal F1 descendants (MP and PM), the parental genome was reciprocally ‘primed’ by *Pst* infection. Although improved defense response against *Pst* infection was observed in both descendants, MP showed a higher resistance when compared to PM (Fig. 1B). The results above showed that the main difference between the MP and PM is CHH methylation. We predicted that CHH DMRs between MP and PM play a role in contributing to their difference in disease resistance and responses. To maximize the identification of potential CHH DMRs, the absolute methylation difference of CHH was set to 0.1. As a result, we identified 761 CHH-hypermethylated DMRs and 285 CHH-hypomethylated DMRs in the MP relative to PM (PM - MP) (Table S3). Methylation levels of most CHH DMRs were higher in MP than that in PM (Fig. 6A; *P* = 3.535e^-09^, Wilcoxon rank-sum test), indicating that MP show CHH hypermethylation, and it is consistent with the previous results. According to the distribution of CHH DMRs in genome elements (Fig. 6B), CHH DMRs are highly enriched in TEs. It is worth noting that in addition to TEs, CHH hyperDMRs are mainly distributed at gene promoter regions.

**Fig. 6.**
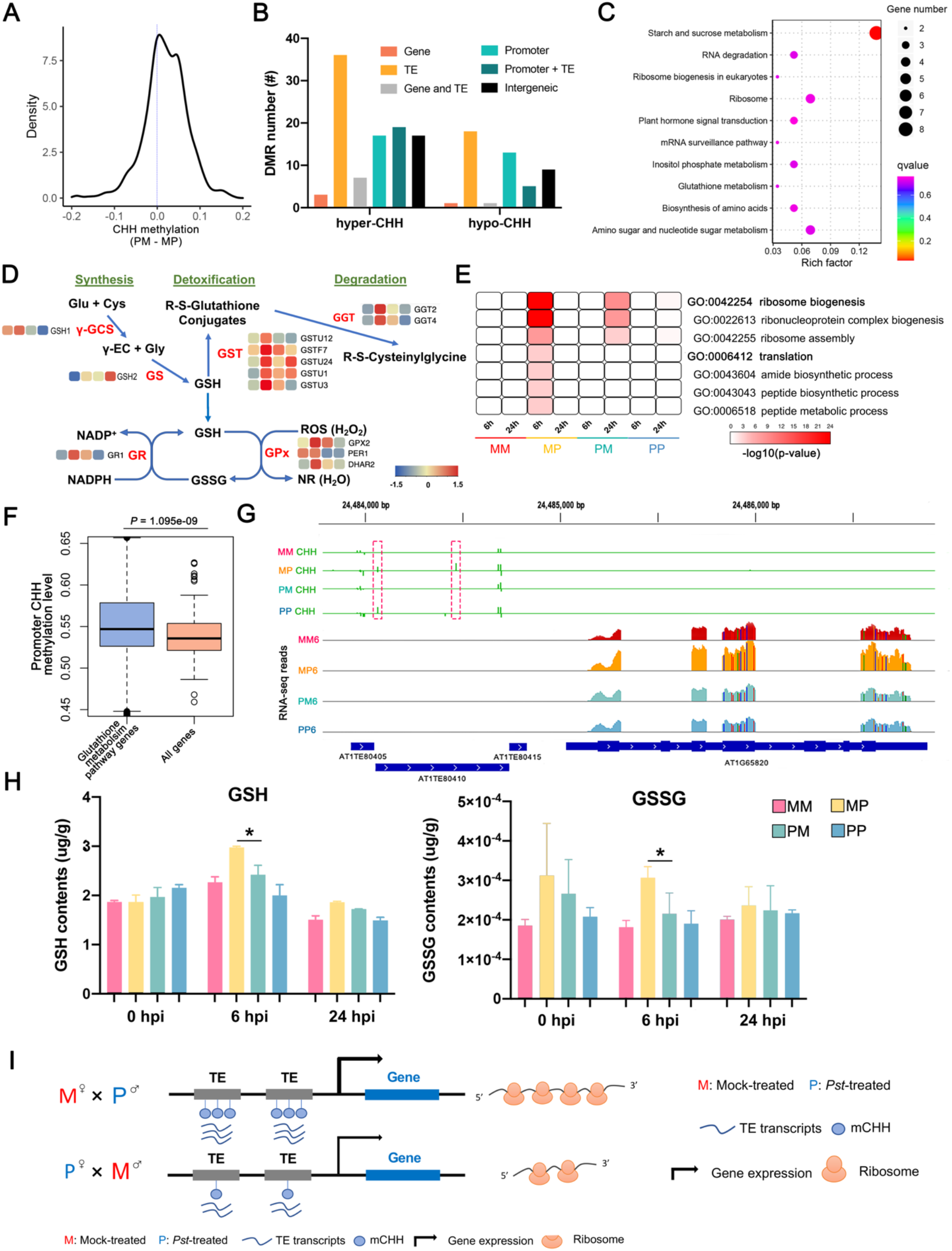
CHH hypermethylation contributes to the disease resistance in MP. (A) Kernel density plot of CHH methylation change of PM/MP in CHH DMRs. (B) The distribution of MP CHH hyperDMRs in genome elements. (C) KEGG analysis of genes whose promoter overlapped with CHH hyperDMRs. (D) Metabolic pathway of glutathione synthesis, degradation and detoxification. Heatmaps showing the RNA expression pattern of genes in pathway were indicated. Color blocks indicate Z-score after normalization, from left to right are MM, MP, PM and PP at 6 hours after *Pst* infection, respectively. γ-GCS, γ-glutamylcysteine synthetases; GS, glutathione synthetases; GST, glutathione S-transferases; GR, glutathione reductases; GPx, glutathione peroxidases. (E) Heatmaps showing the translation-related GO terms enriched with upregulated genes across F1 descendants. (F) Boxplot showing CHH methylation of promoter in glutathione metabolism genes and all annotated protein-coding genes in *Arabidopsis* genome. (*P* value < 0.001, as determined using the t-test). (G) IGV view of the AT1G65820 promoter CHH methylation levels and RNA-Seq reads across the MM, MP, PM and PP at 6 hours after *Pst* treatment. Regions of large differences in CHH methylation are highlighted by the dotted outline of a rectangle. Gene models are represented in blue. (I) GSH and GSSG contents in MP and PM treated with *Pst* at the 0, 6 and 24 hours. Asterisks represent significant difference at *P*-value < 0.05 in t-test. (H) Schematic diagram illustrating CHH methylation in TE contributing to altered transcriptome changes, especially genes involved in ribosome biogenesis in reciprocal descendants of *Pst-*primed parents.

It has been reported that DNA methylation can target promoter region to regulate stress responsive genes (Le et al., 2014). The presence of TEs in the vicinity of genes has been reported to associated with their imprinted status (Rodrigues and Zilberman, 2015). To gain insights into the functional consequence of DMRs between MP and PM, we used KEGG database to categorize genes whose promoters are overlapped with CHH hyperDMRs. KEGG pathway of ‘Starch and sucrose metabolism’, ‘Ribosome’ and ‘Glutathione metabolism’ (Fig. 6C) were significantly enriched among genes exhibiting CHH hypermethylation at promoter regions. Interesting, consistent with the transcriptomic level (Fig. 5D), at methylation level, the glutathione metabolism was also enriched in MP. Most of these KEGG pathway have been reported to be related to stress responses. For example, the starch and sucrose is engaged in plant defense by activating plant immune responses against pathogens (Tauzin and Giardina, 2014). In addition, it has been shown that glutathione is required for *Arabidopsis* immunity (Ball et al., 2004; Hiruma et al., 2013). Moreover, pathways related to translation process, such as ‘Ribosome’, ‘Biosynthesis of amino acids’ and ‘Ribosome biogenesis in eukaryotes’ are also enriched, which is a fundamental layer of immune regulation in plants (Xu et al., 2017). Collectively, the CHH methylation showed correlation with stress responsive genes regulation in MP.

At both methylation and transcriptomic level, the glutathione metabolism KEGG pathway was enriched in MP. We found that transcripts related to glutathione synthesis, detoxification and degradation was upregulated in MP6 when compared with other samples (Fig. 6D). For example, a glutathione detoxification-associated genes glutathione S-transferase 7 (GSTF7) and four glutathione S-transferase TAU genes (GSTU1, 3, 12 and 24) were all induced in MP. In plants, glutathione participates in detoxification as well as signaling in defense against the pathogens (Ghanta and Chattopadhyay, 2011), indicating the potential of altered glutathione metabolism in contributing to the disease resistance in MP6. γ-glutamylcysteine synthetase (γ-GCS), one of the glutathione synthesis enzymes, was upregulated in MP after *Pst* infection. Enzymes related to glutathione scavenging reactive oxygen species (ROS) are also upregulated in MP and play a potentially protective role in stress (Fig. 6D). Previously study has been showed that increased glutathione contributes to stress tolerance and global translation changes in *Arabidopsis* (Cheng et al., 2015). Thus, we then compared the GO analysis of DEGs between the untreated (0 hours) and *Pst*-treated (6 and 24 hours) in F1 descendants. Genes involved in processes associated with the ribosome biogenesis and translation GO terms (Fig. 6E, Fig. S9) are overrepresented among the up-regulated genes in MP. The translation GO terms (Fig. S9) and core histone (CH) gene family (Fig. S12) are also enriched in genes which were upregulated in reciprocal descendants, which were more significantly in MP.

Besides, we found that the CHH methylation level of the promoters of glutathione metabolism pathway genes is significantly higher than that of all protein-coding genes (Fig. 6F). For example, a microsomal glutathione s-transferase (AT1G65820), which containing three TEs in its promoter, showed higher CHH methylation in MP when compared to PM and was upregulated in MP after *Pst* infection (Fig. 6G). To further test whether CHH methylation promotes the accumulation of glutathione, we measured the contents of GSH and GSSG in the MP and PM at 0, 6 and 24 hours. After the *Pst* treatment, the GSH and GSSG levels in MP were significantly higher than PM at 6 hours after *Pst* treatment (Fig. 6H). Taken together, the paternal contribution could lead the CHH hypermethylation, potentially through glutathione to enhance gene expression and translation for enhance disease resistance in MP (Fig. 6I).

## Discussion

Although parental stress effects are well documented in plants, the extent to which molecular phenotypes are affected by stress exposures in previous generation is not well understood. Plants use a series of defense mechanisms to restrict the growth of biotrophic bacteria upon infection, such as defense gene expression, which is modulated by DNA methylation. In plants, DNA methylation and histone acetylation have emerged as critical regulators of defense priming. DNA methylation is dynamic but also incredibly stable between generations. Difference in DNA methylation is a potential source of epialleles that can lead to phenotypic diversity, and this could be captured or created for crop improvement (Springer and Schmitz, 2017). Research suggests that although most methylation loci are stably inherited, locations of epialleles have varying stability over generational time. Very few multi-omics approaches have been performed to the effects after parental exposure to cues associated with pathogen. So, it important to understand the parental contribution of epigenetic variation and fully utilize the potential of epigenetic variation for crop improvement.

In this study, all descendants with one or more primed genome (MP, PM, and PP) showed higher disease resistance than unprimed genome descendant (MM), suggesting a unique parental contribution for improved stress responses in the progenies. In addition, many histone-related genes were upregulated in the primed genome, suggesting a potential of chromatin organization rearrangement (Fig. 5F). The main difference between the unprimed (MM) and primed (PP) genome is CG methylation (Fig. 3A), which can be stably inherited to the next generation (Stassen et al., 2018). Stress-primed genome induces regulatory changes at both paternal and maternal alleles, causing significant changes in methylation level and gene expression in the F1 descendants. The differences in methylation state may explain changes in transcriptome among different descendants. The primed genome are rapidly and strongly induced after *Pst* infection, which is contributed indirectly by more rapid changes in chromatin organization, leading to a more rapid response (defense-related gene expression changes) against *Pst* infection. PP mainly enhances its own disease resistance ability by rapidly upregulating the disease resistance genes (Fig. 4) and increasing photosynthesis to delayed precocious senescence (the chlorophyll a/b-binding gene family were significantly overrepresented in PP when compared to MM (Fig. S12). For reciprocally primed F1 descendants (MP and PM), epigenome confrontation triggers reprogramming of DNA methylation, leading to transgressive segregation accompanied with chromatin changes and enhanced gene expression involved in translation process, immune response and defense response (Fig. S8; Fig. S11). In this study, the *Tc1* transposable element family were highly expressed in reciprocal F1 descendants. It has been reported that *Tc1* shapes genetic variation and gene expression in the protist *Trichomonas vaginalis* (Bradic *et al.*, 2014). Therefore, it is possible that the *Tc1* family may form new epialleles, contributing to transgressive segregation in reciprocal descendants.

One noteworthy aspect of this study is that self-fertilized F1 descendants (MM and PP) and reciprocally crossed progenies (MP and PM) were generated differently. MM and PP were from natural self-pollination without the artificial emasculation and pollination process. Artificial pollination cannot avoid affecting the vitality of pollen, and then affect some traits of descendants (Hopping and Hacking, 1982; Richardson and Anderson, 1996). In this study, we focus on exploring the influence of paternal contribution and maternal contribution on disease resistance of descendants, and artificial pollination (Figure 4) and self-pollination (Figure 5 and 6) samples were compared separately to reduce the impact of artificial pollination as much as possible. Therefore, we believe that different pollination methods have little effect on the conclusions of this study.

Primed paternal genome (MP) induces CHH hypermethylation in the genomic loci with TEs and stress-responsive genes, establishing a ‘priming’ state that potentiates transcriptional regulation upon stress. Some TEs could activate nearby genes, while most of TEs are transcriptionally silenced in plant genome (Lin-Wang et al., 2010; Lippman et al., 2004). In maize, the CHH methylation exhibited a positive correlation with gene expression (Gent et al., 2013). Transcription initiates from TEs through Pol IV or Pol II and can spreads to nearby genes (Haag and Pikaard, 2011; Zheng et al., 2009). Parental CHH methylation was also reported to affect circadian rhythms and biomass heterosis in *Arabidopsis* intraspecific hybrids (Ng et al., 2014). In our study, CHH hypermethylation were accompanied by glutathione pathway and enhancement of gene expression related to translation. The effect that we observed on offspring after *Pst* infection is consistent with function related to the pathogen resistance, with both methylomes and transcriptomics analysis converging on glutathione signaling as a paternal affected pathway. Therefore, our results suggested that paternal contribution promote *de novo* CHH methylation at promoter regions of disease resistance genes (also including TEs) and potentially contribute to more disease resistance in MP in response to *Pst* challenge.

From our transcriptome analysis, we found that pathway related to SAR, JA (Fig. S11) and glutathione metabolism (Fig. 6D; Fig. S11) were overrepresented in MP, suggesting paternal priming by *Pst* can strengthen these pathway genes for enhanced disease resistance in F1. Increased JA levels and altered jasmonate signaling can mediate long-distance information transmission during SAR. Interestingly, JA treatment in *Arabidopsis* was found to induce expression of glutathione metabolic genes, thereby improving the capacity of synthesizing and recycling of glutathione, potentially rendering the plants to be more responsive when subjected to stress (Xiang and Oliver, 1998). In contrast, the GO terms related to stress (including defense response, defense response to bacterium and response to biotic stimulus) were more enriched in PM than MP (Fig. S11), when compared with PP. Besides, the glutamate receptor (GR) gene family (Fig. S12) are also more enriched in PM, and the GR genes have been implicated in plant defenses to biotic stress. Our analysis indicates that the parental effects of the two different epigenetic genome were differentially regulated. These findings may provide additional sources of epigenetic variations within a species that could be captured or created for disease resistance, and facilitate further investigation of parental contributions to enhance our understanding of the molecular basis of disease resistance for breeding selection.

## Methods

### Establishment of stress-primed progeny lines

Parental plants (L*er*) grown under short-day conditions (8-h light/ 16-h dark) at 22°C to delay bolting. Prior to bolting, 5-7 weeks old mature plants were subjected to 4 consecutive treatments with *Pseudomonas syringae* pv. *tomato* DC3000 (*Pst*DC3000) at intervals of 3-4 days by dipping inoculation. For priming, an inoculum of 1×10^8^ CFU/ml was used for the 1^st^ and 2^nd^ treatments and an inoculum of 1×10^9^ for the 3^rd^ and 4^th^ treatments, respectively. Parallel mock treatments with 10mM MgSO4, 0.01% Silwet L-77 was included. At 3 weeks after the final treatment, emasculation and pollination were performed when there are sufficient numbers of the inflorescence. The self-fertilized F1 descendants (MM and PP) were generated from natural self-pollination without the artificial emasculation and pollination. In total, we obtained about 3∼6 crosses plant for each progeny line. F1 descendants were collected for subsequent characterization of their defense responses against *Pst*DC3000.

### Plants treated with *Pst*DC3000 in F1 descendants and sampling

For F1 descendants, plants are grown under short-day conditions (8-h light/ 16-h dark) at 22°C. Then 5-week-old plants were subjected to mock-, *Pst*DC3000 treatment in parallel by syringe infiltration. Their disease resistance was assessed based on the development of disease symptoms (chlorosis and necrosis) among different lines at 0, 2 and 3 days post infection (dpi). In addition, the level of disease resistance was scored and the bacterial population in the inoculated leaf tissues was determined and quantified as colony forming units (CFU) per leaf area. For WGBS-seq, samples were collected at 0 hours post treatment (hpt). For RNA-seq, samples were collected at 0, 6 and 24 hpt. In total, three biological replicates were performed for the experiment. Each replicate sample for each data point was collected from three plants.

### Whole-genome bisulfite sequencing and DMR analyses

For each descendant, genomic DNA was extracted using QIAamp DNA Mini Kit (QIAGEN) from 5-week-old leaves of MM, MP, PM and PP lines. We did three biological replicates per descendant. Whole-genome bisulfite sequencing were performed at Beijing Genomics Institute (BGI) using HiSeq2500 platform, producing 100-bp paired-end reads (Supplemental Table S1). Read quality was assessed with FastQC and trimmed using FASTX-Toolkit (http://hannonlab.cshl.edu/fastx_toolkit/). Clean reads were mapped to TAIR10 genome using the Bismark (v0.22.3) (Krueger and Andrews, 2011) allowing two mismatches. Bases covered by fewer than 5 reads were excluded, and only uniquely mapped reads were used for further analysis. The efficiency of bisulfite conversion was calculated by aligning the reads to the lambda genome (which is fully unmethylated). Whole-genome bisulfite sequencing statistics are provided in Table S1.

To identify regions of the DMRs between two descendants, DMRs were defined by comparing the methylation level of 200bp windows throughout the genome between two samples using the methylkit (Akalin et al., 2012). Bins with false discovery rate (FDR) < 0.01 and absolute methylation difference of 0.4, 0.2, 0.2 for CG, CHG, CHH were defined as hypermethylation/hypomethylation, respectively. To avoid 100-bp bins with few cytosines, we selected bins with at least four cytosines that are each covered by at least ten reads and maximum of 100 reads in each sample. Finally, DMRs within 100 bp of each other were merged by allowing a gap of one window.

### RNA-Seq data processing

The tissues were collected and frozen immediately in liquid N2, and total RNA was extracted using the Purelink Plant RNA Reagent according to the manufacturer’s protocol. mRNA sequencing was outsourced to Beijing Genomics Institute (BGI), and libraries were sequenced on BGISEQ-500 platform. In briefly, DNase I was initially used to degrade DNA contaminant in RNA samples. The mRNA molecules were then purified from total RNA using oligo(dT)-attached magnetic beads and fragmented into small pieces. First-strand cDNA was generated by reverse transcription PCR using random hexamer primers, followed by a second-strand cDNA synthesis. Subsequently, A-Tailing Mix and RNA Index Adapters were added to perform end-repair. The double-stranded PCR products were heat-denatured together and circularized by the splint oligo sequence to obtain the final library. The single-stranded circular DNA was amplified using phi29 to generate DNA nanoball. The DNBs were loaded into the patterned nanoarray and single-end 50 bp reads were generated on the BGISEQ-500 sequencing platform (Huang et al., 2017). Three independent biological samples were sequenced per sample. Raw data were quality- and adaptor-trimmed using Fastx-Tookit. Next, all clean reads were mapped independently to the *A. thaliana* genome using the program STAR v2.7.3a (Dobin et al., 2013) with default parameters (Table S2). The number of reads mapped to each annotated gene were determined by FeatureCount v1.6.4 (Liao et al., 2013). Araport11 gene and TE models were obtained from TAIR (www.arabidopsis.org). Finally, raw read counts were normalized to homoscedastic expression values using *variance stabilizing transformation* (VST) function implemented in the DESeq2 package (Love et al., 2014). Principal component analysis (PCA) was performed used the DESeq2 package (Love *et al.*, 2014) based on the transformed data (Fig. S3). The number of aligned reads for each gene was normalized to TPM (Transcripts Per Million), which was used to represent the gene and TE expression level.

DESeq2 Bioconductor package (Love *et al.*, 2014) was used to identify differentially expressed genes (DEGs). DEGs were calculated among the three time points (e.g., 0h vs. 6h; 0h vs. 24h; 6 vs. 24h). For each pair of samples, we compared the resulting read counts from three biological replicates. For each time point, we performed pairwise comparisons among F1 descendants, including MM vs. MP, MM vs. PM, MM vs. PP, MP vs. PM, MP vs. PP and PM vs. PP. The DEGs were determined based on |log2-fold change| > 1 and FDR of < 0.01.

### GO term, KEGG and gene family analysis

The GO term analysis was performed using the web-based tool AgriGO v2 with singular enrichment analysis (Tian et al., 2017), and GO terms with FDR < 0.05 were identified as significant terms. Then REViGO (Supek et al., 2011) was used to generate GO term redundancy clusters. KEGG enrichment analysis was performed with the R package clusterProfiler (Yu et al., 2012), with a Bonferroni correction and an adjusted *p*-value of 1. The gene family enrichment analysis was performed on GenFam website (Bedre and Mandadi, 2019) with FDR < 0.05. TAIR10 genome was used as background.

### Quantification of GSH and GSSG contents

The GSH and GSSG contents were measured using the Reduced Glutathione (GSH) Content Assay Kit (Sangon Biotech, Shanghai, China, NO. D799614) and Oxidized Glutathione (GSSG) Assay Kit (Sangon Biotech, Shanghai, China, NO. D799616) following manufacturer’s instructions, respectively.

## Data availability

All WGBS-seq and RNA-Seq data from this study are available from the SRA database under the accession number PRJNA718691.

## Author contributions

YPL, DWKN designed the research; EYML, XSZ, TJL performed the research and collected data; YPL and DWKN analyzed and interpreted data; YPL wrote the manuscript with input and edits from DWKN.

## Acknowledgements

We thank Yee Ching Ng for her technical help during the early stage of the project.

---

**Table 1.**
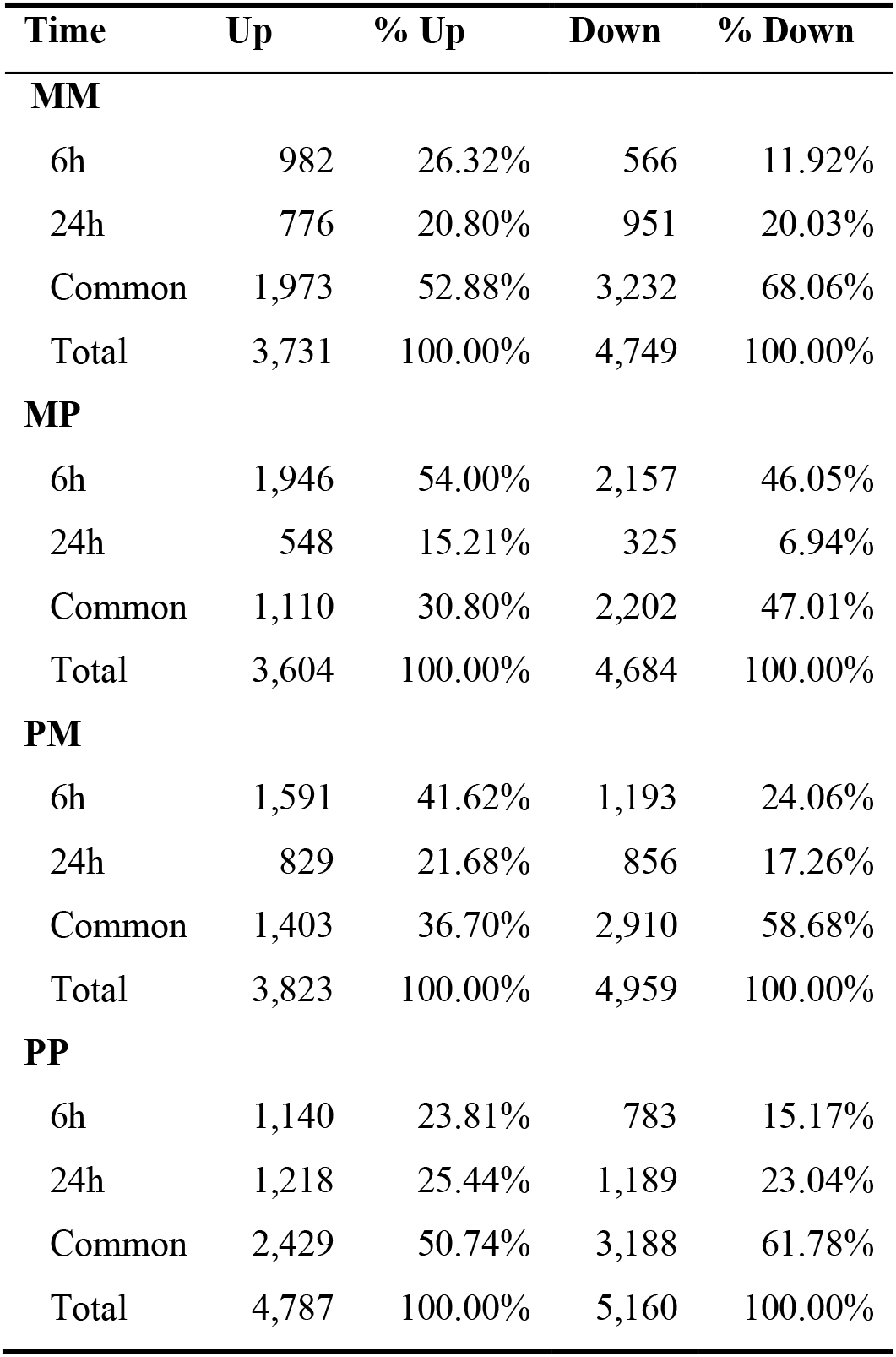
Proportion of up- and downregulated genes in response to *Pst* infection.

## Supplemental Information

**Figure S1.**
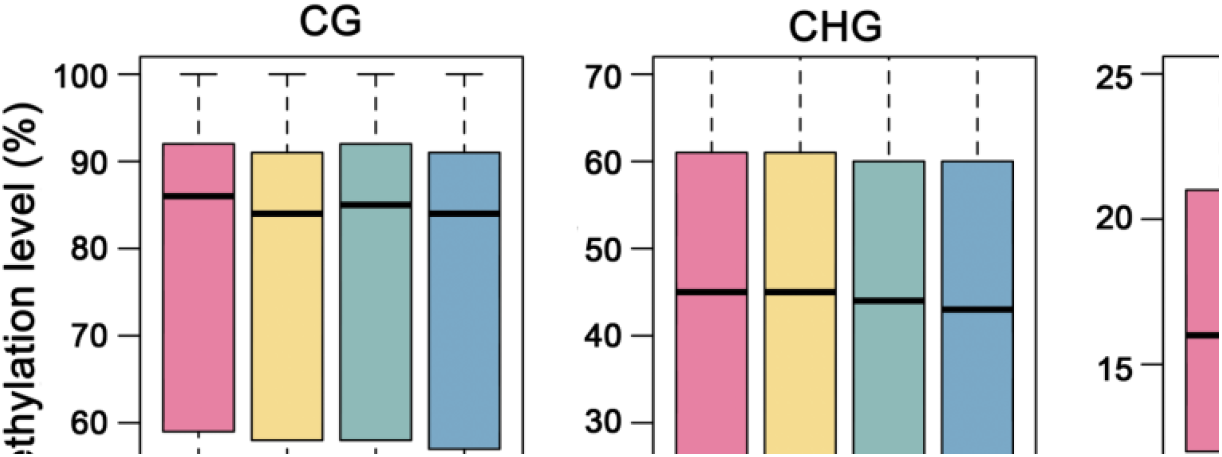
Comparison of mean methylation content in MM, MP, PM and PP. *Arabidopsis* genome was divided into 100bp bins, boxplots showing the average methylation level of each bin. The y-axis shows the average methylation level of MM, MP, PM and PP in CG, CHG and CHH contexts, only cytosines covered by at least five reads were considered. The methylation levels were derived from three biological replicates for each genotype.

**Figure S2.**
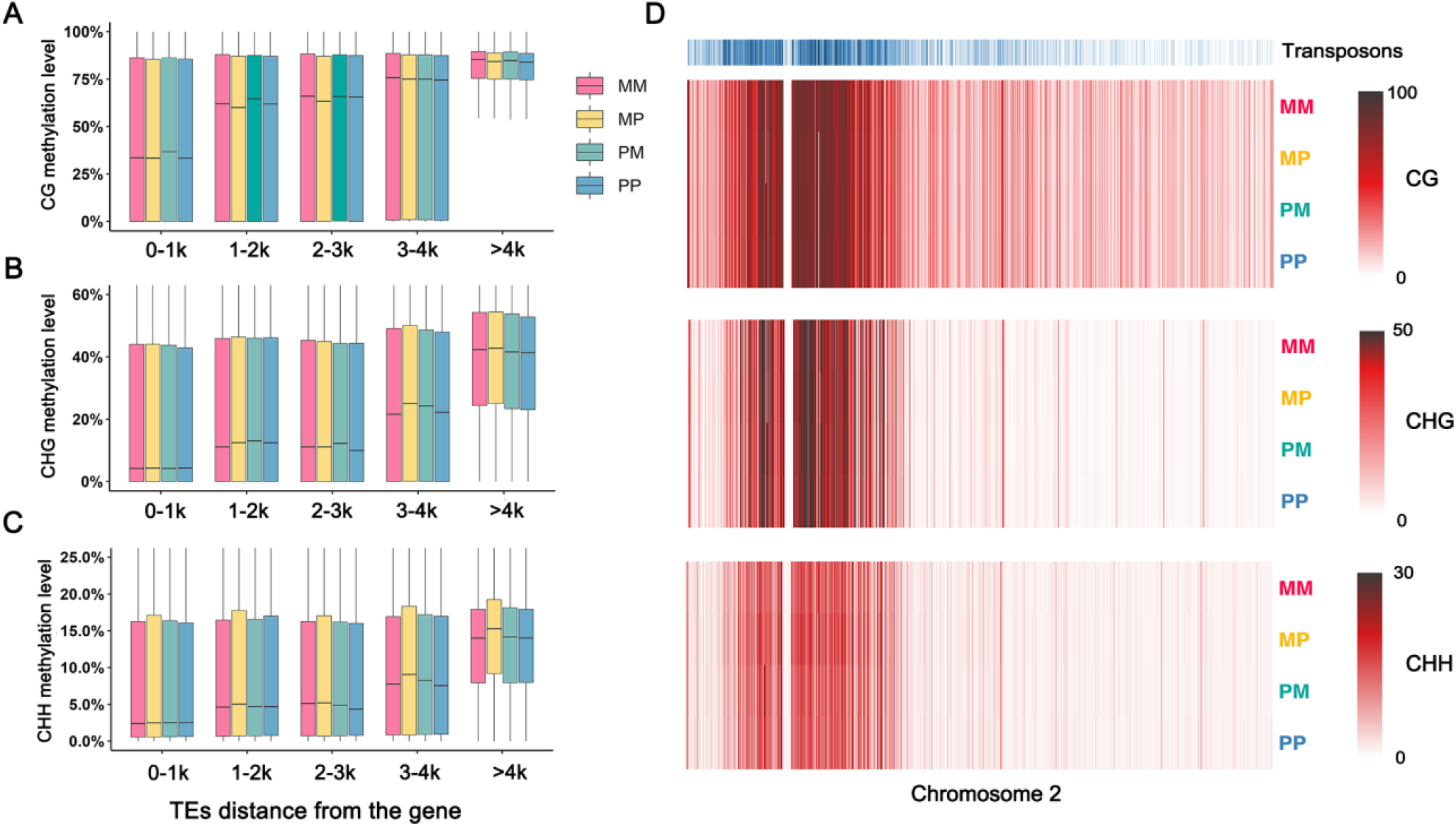
MP CHH hypermethylation is not restricted to the pericentromere. (A-C) TE methylation level of CG, CHG and CHH contexts relative to the distance from the nearest gene in F1 descendants. (D) Heatmaps of transposon density or methylation level in 20-kb windows across chromosome 2. Each methylation context has its own scale bar to visualize changes across different descendants.

**Figure S3.**
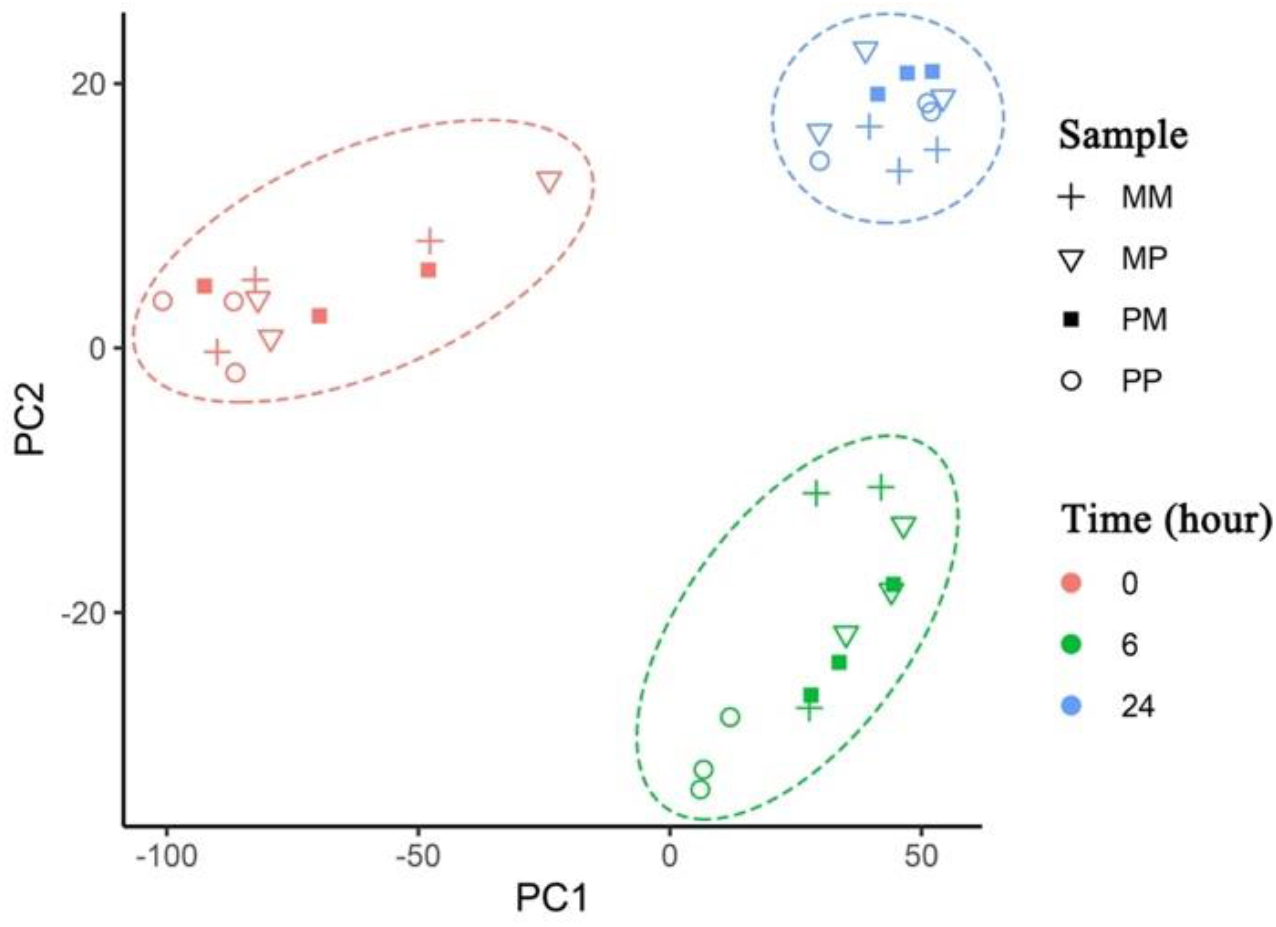
Principal component analysis (PCA) of gene expression levels in *Arabidopsis thaliana* plants treated with *Pst*DC3000. PCA was executed with DESEQ2 software on the *variance stabilizing transformation* (VST) data. The first two principal components are plotted for data from F1 descendants, including MM (+), MP (s), PM (▪), and PP (O), at different time points (0, 6, and 24 hours) after challenged with *Pst*DC3000 (2 x 10^8^ cfu/ml, OD600 = 0.4). Percentages of variation explained by each PC are indicated along the axes. MM and PP are direct descendants from the mock-treated and *Pst*-treated parents, respectively. MP and PM are F1 descendants from reciprocal crosses between a mock-treated and a *Pst*-treated parent.

**Figure S4.**
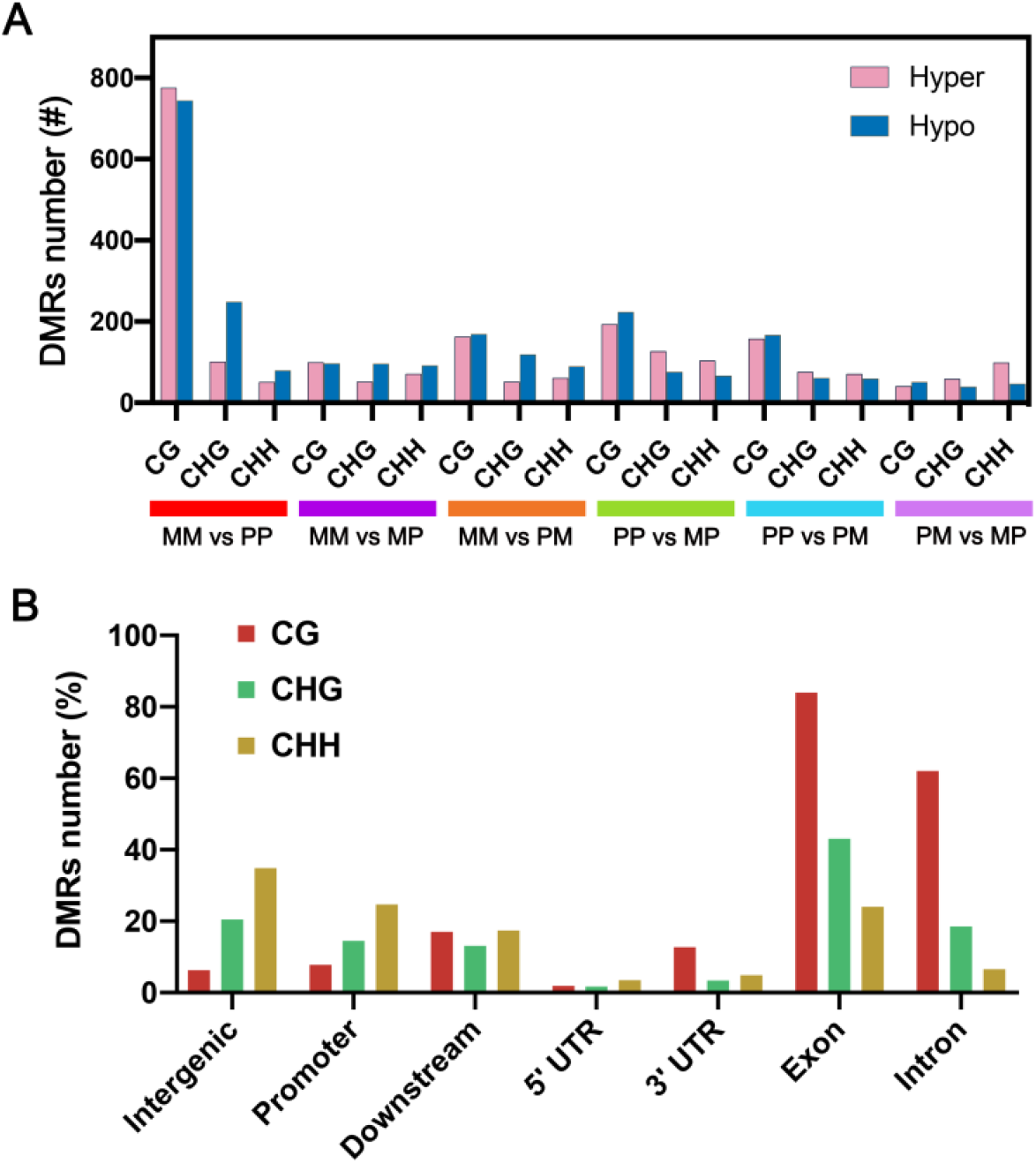
DMRs and their distributions among F1 descendants. (A) Summation of all DMRs genome-wide across individual genotypes. Absolute methylation differences of +/- 40% for CG, +/- 20% of CHG and CHH were defined as hypermethylation/hypomethylation, respectively. (B) Distribution of identified all merge DMRs in genomic features. Promoter and downstream regions were defined as 1 kb upstream of the transcription start site and 1 kb downstream of the transcription termination site, respectively.

**Figure S5.**
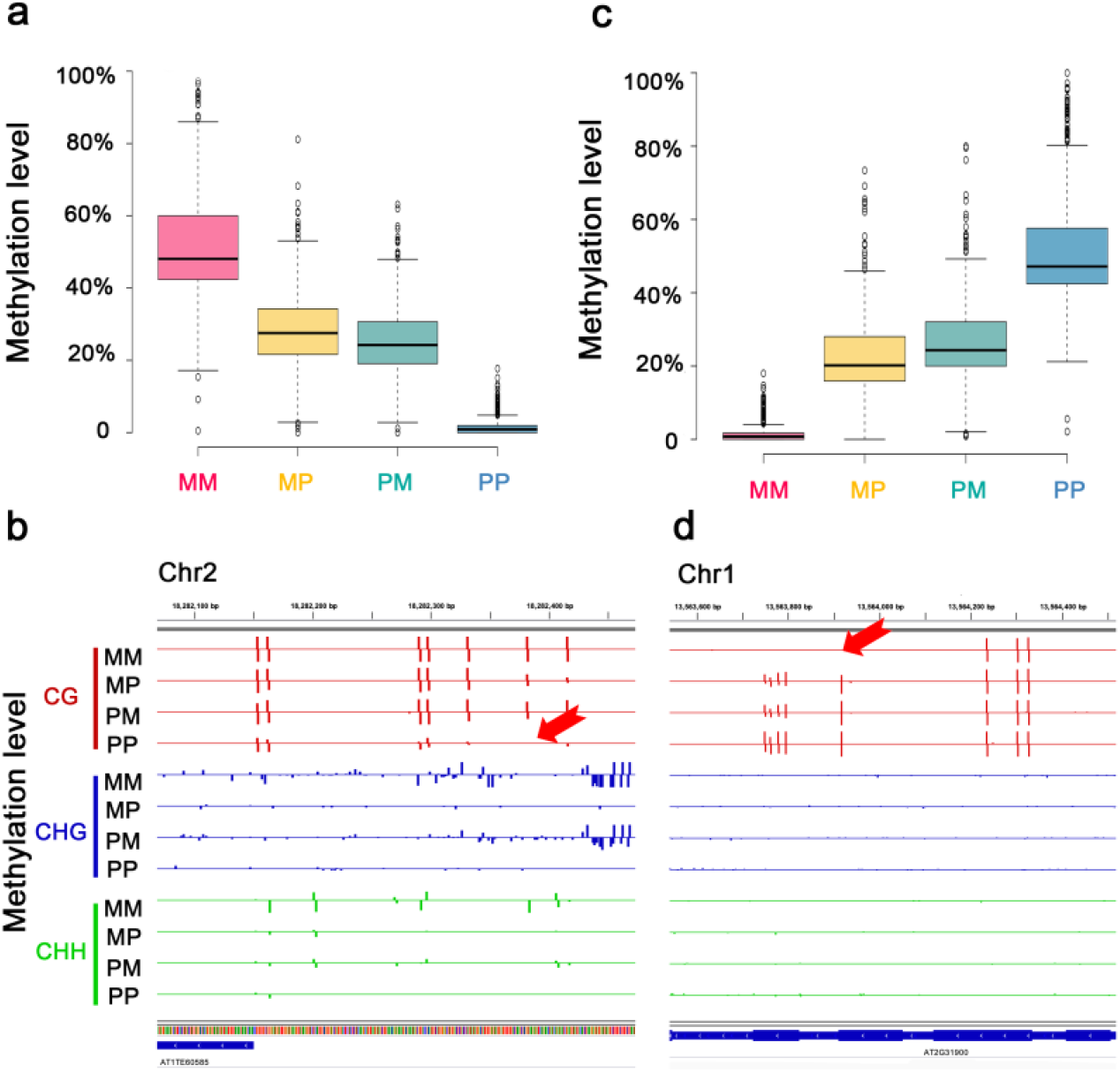
Methylation analyses of regions that are differentially methylated in MM and PP. (A) Boxplot showing methylation levels of 364 regions highly methylated only in MM. (B) A region adjacent to *AT1TE60585* that is highly methylated in MM. (C) Boxplot showing methylation levels of 407 regions highly methylated only in PP. (D) A genic region (*AT2G31900*) that is highly methylated in PP.

**Figure S6.**
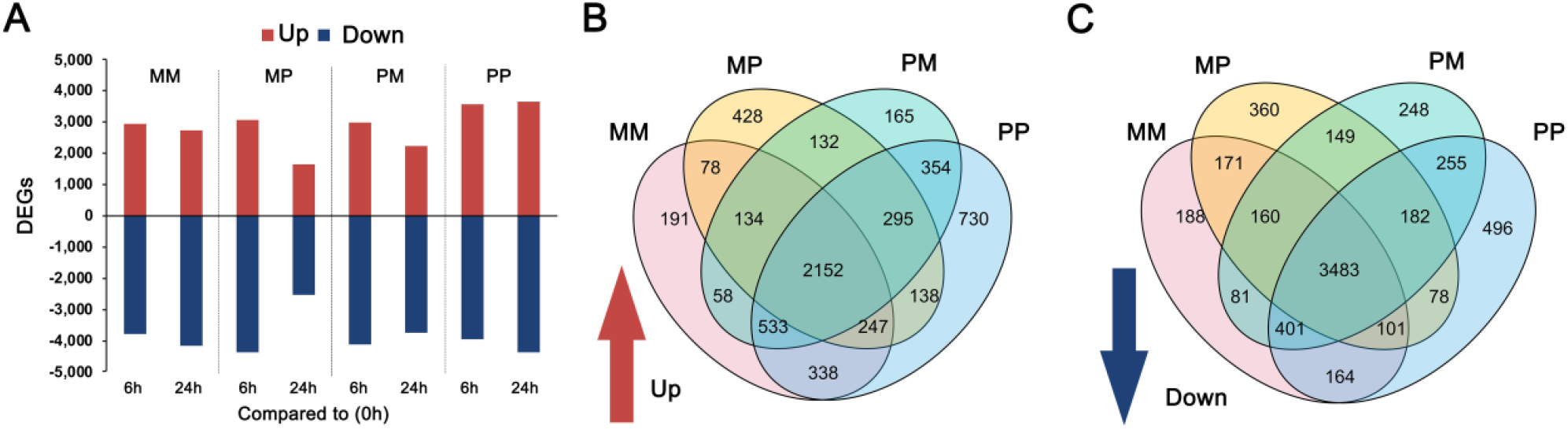
Transcriptome analysis of F1 descendants exposed to *Pst*DC3000. (A) The number of differentially expressed genes (DEGs) at 6 and 24 hours after *Pst* treatment. Genes were considered to be differentially regulated if they displayed a log 2-fold change ⩾1 (upregulated) or ⩽ –1 (downregulated). p-value was adjusted for multiple testing using *Benjamini-Hochberg* method (padj) ⩽ 0.01. (B-C) Venn diagrams showing number of (B) up- and (C) down-regulated DEGs in response to *Pst* challenge.

**Figure S7.**
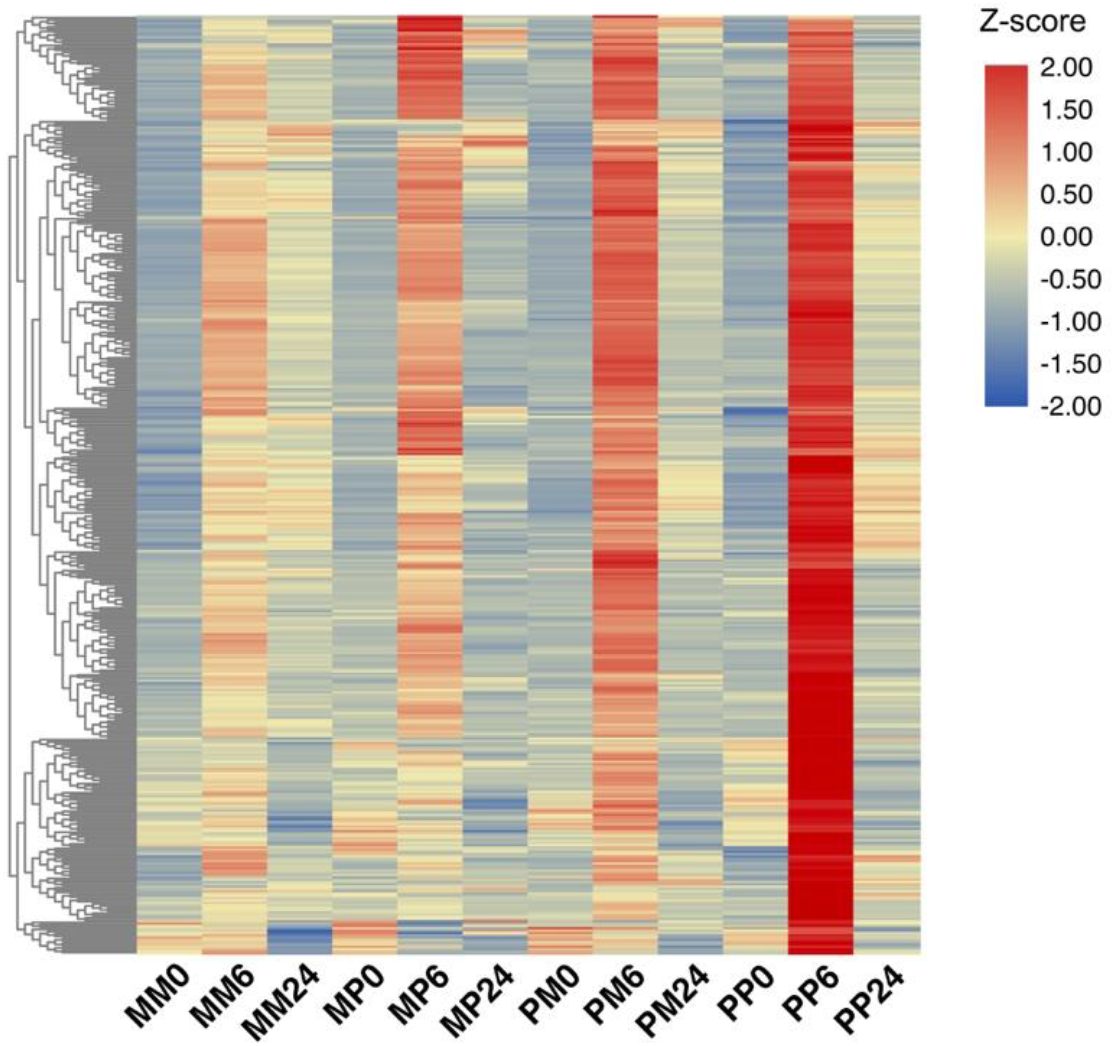
Expression profiles of identified primed genes. Heatmap showing the expression profiles of the identified differentially expressed genes after *Pst* treatment among the F1 descendants at the indicated time point. Z-score obtained from averaged TPM of three biological replicates.

**Fig. S8.**
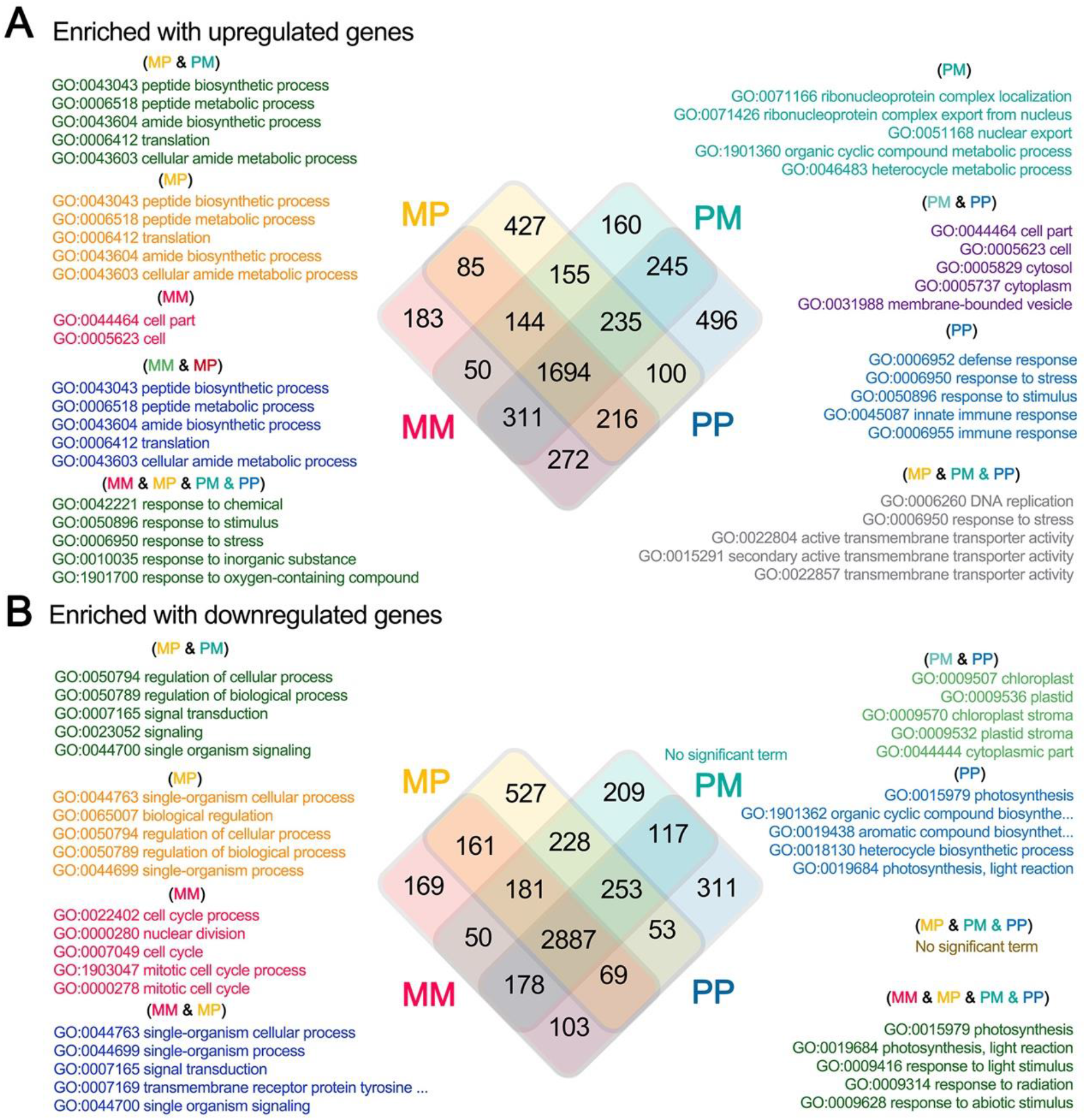
Distribution of differentially expressed genes (0 vs. 6 hours) according to gene ontologies (GOs). Distribution of GOs enriched with upregulated genes (A) and downregulated genes (B) across F1 descendants. MM and PP are direct descendants from the mock-treated and *Pst*-treated parents, respectively. MP and PM are F1 descendants from reciprocal crosses between a mock-treated and a *Pst*-treated parent.

**Figure S9.**
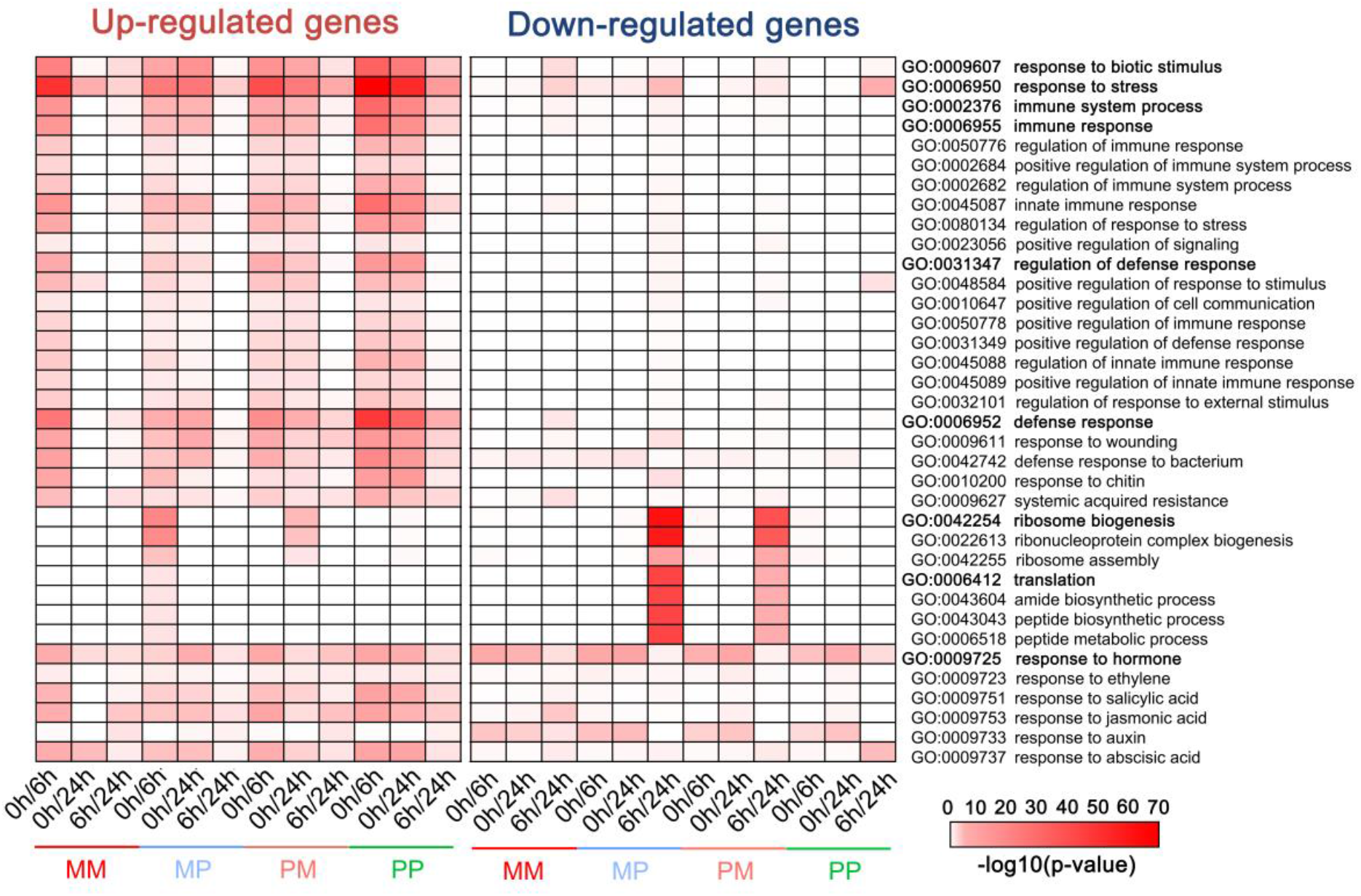
Gene Ontology enrichment of differential expressed genes. Prominent GO terms enriched in the up- and down-regulated DEGs of the MM, MP, PM and PP exposed to *Pst*DC3000 over time. (P ⩽ 0.01; for full GO listing, see Supplement Dataset S3). The intensity of the red color bar indicates p-value values on a –log10 scale.

**Figure S10.**
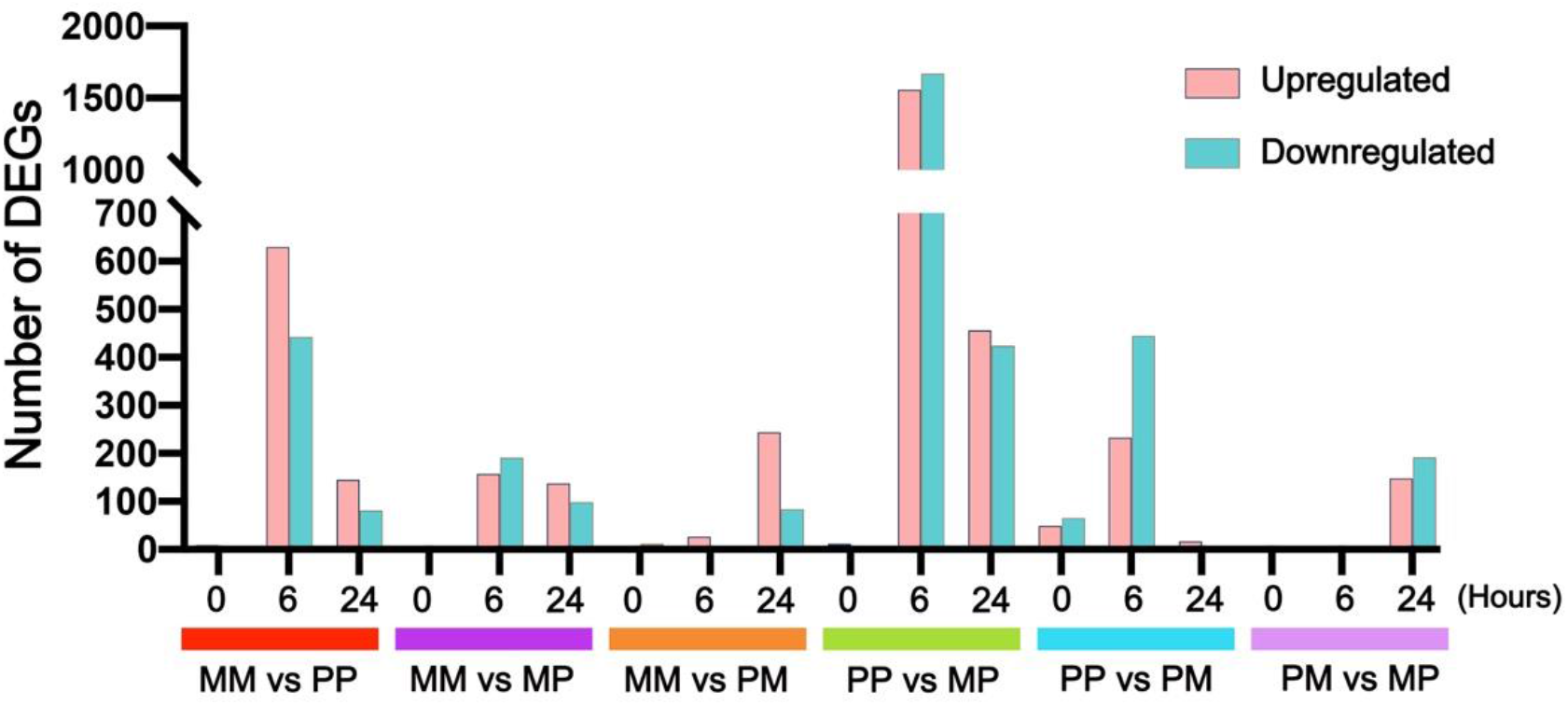
Number of DEGs between different F1 descendants in response to *Pst* treatment. Bar plot shows the number of differentially expression genes (DEGs) in each combination at each time point after *Pst*DC3000 treatment.

**Figure S11.**
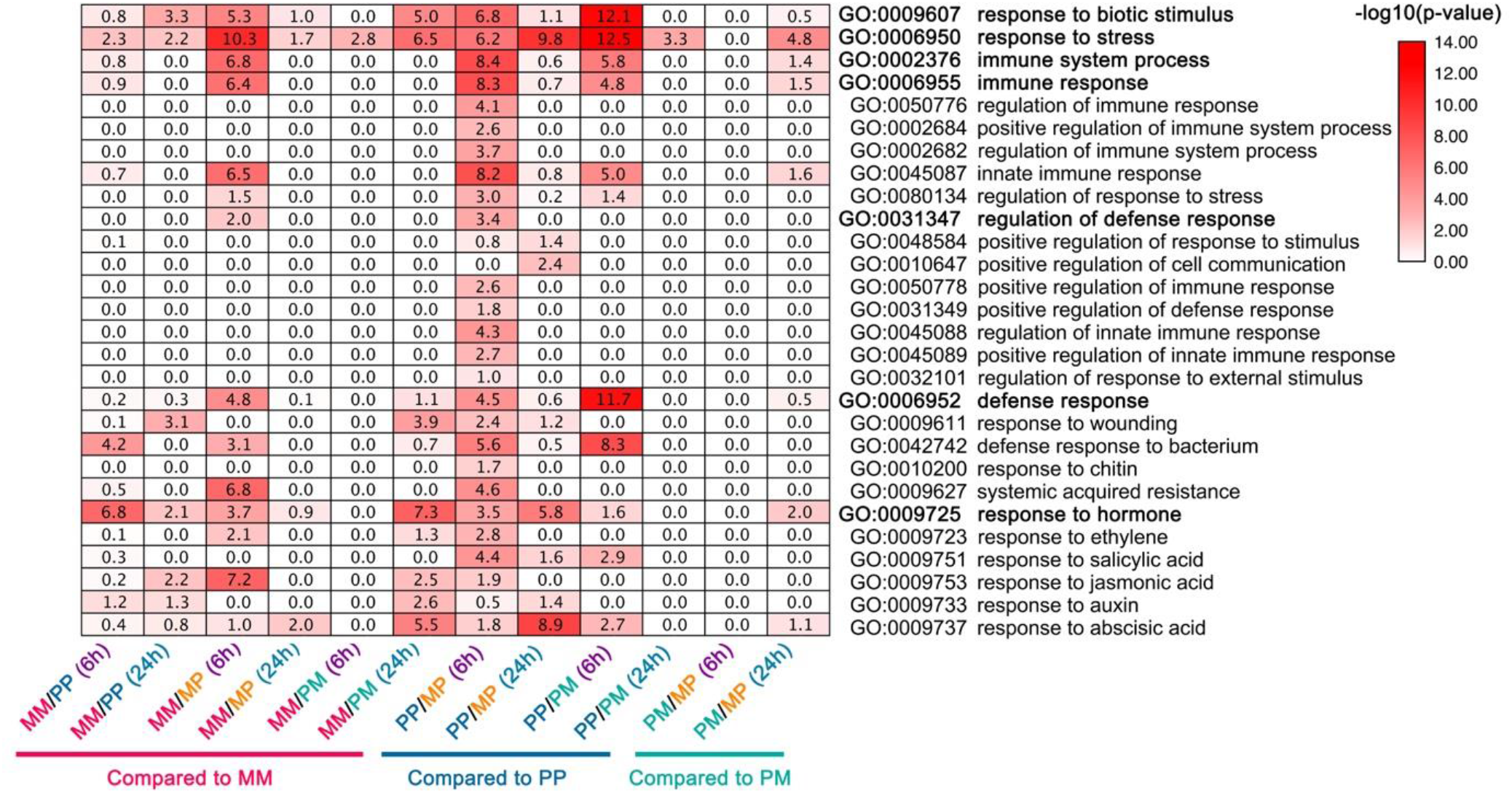
Up-regulated DEGs Gene Ontology. Prominent GO terms enriched in the up-regulated DEGs in pairwise comparisons (MM vs. PP, MM vs. MP, MM vs. PM, PP vs. MP, PP vs. PM, PM vs. MP) when exposed to *Pst*DC3000 over time.

**Figure S12.**
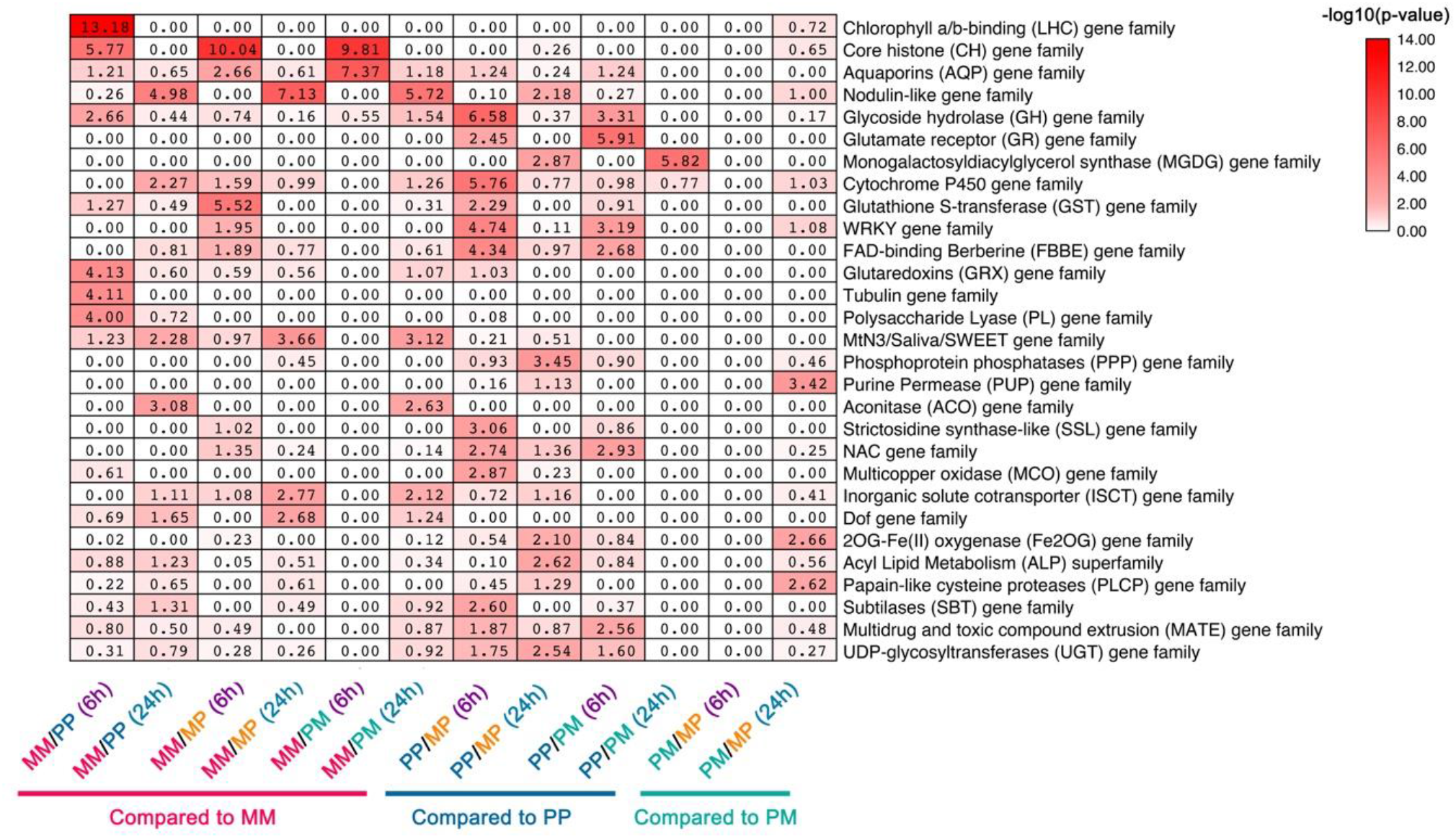
Gene families of up-regulated DEGs. Prominent gene family enriched in the up-regulated DEGs in pairwise comparisons (MM vs. PP, MM vs. MP, MM vs. PM, PP vs. MP, PP vs. PM, PM vs. MP) when exposed to *Pst*DC3000 over time (P ⩽ 0.01). The intensity of red color bar indicates a –log10 scale of the p-value.

**Table S1.** Whole-genome bisulfite sequencing statistics.

| Samples | Number of clean paired-end Reads | Number of mapped paired-end reads | Coverage | Total methylated C's in CpG context (% methylated) | Total methylated C's in CHG context (% methylated) | Total methylated C's in CHH context (% methylated) | Bisulfite conversion rates (%) |  |  |
| --- | --- | --- | --- | --- | --- | --- | --- | --- | --- |
|  |  |  |  |  |  |  | CG | CHG | CHH |
| MM0_1 | 24,524,728 | 14,044,685 | 23 | 19,214,960 (23.2) | 6,593,937 (7.8) | 11,912,393 (3.3) | 99.4% | 99.2% | 99.1% |
| MM0_2 | 24,742,192 | 14,967,660 | 25 | 20,684,888 (23.2) | 6,905,082 (7.6) | 12,582,836 (3.2) | 99.5% | 99.4% | 99.3% |
| MM0_3 | 24,690,815 | 14,681,812 | 25 | 21,298,076 (24.2) | 7,202,111 (8.0) | 13,426,827 (3.5) | 99.5% | 99.4% | 99.3% |
| MP0_1 | 24,674,563 | 14,327,659 | 24 | 19,103,676 (22.8) | 6,309,122 (7.3) | 12,378,013 (3.3) | 99.4% | 99.4% | 99.3% |
| MP0_2 | 24,691,545 | 14,038,866 | 23 | 198,31,989 (24.2) | 6,959,786 (8.3) | 13,472,157 (3.8) | 99.4% | 99.4% | 99.1% |
| MP0_3 | 24,739,461 | 14,727,607 | 25 | 21,720,258 (24.3) | 7,382,081 (8.1) | 13,902,145 (3.7) | 99.4% | 99.4% | 99.3% |
| PM0_1 | 24,697,520 | 14,205,216 | 24 | 19,971,368 (24.0) | 6,706,643 (7.9) | 12,466,930 (3.4) | 99.4% | 99.2% | 99.1% |
| PM0_2 | 24,550,816 | 13,351,332 | 22 | 19,445,046 (24.1) | 6,557,272 (8.0) | 11,867,420 (3.5) | 99.4% | 99.3% | 99.2% |
| PM0_3 | 24,628,377 | 14,358,187 | 24 | 19,930,520 (23.3) | 6,578,203 (7.5) | 12,252,507 (3.3) | 99.7% | 99.3% | 99.3% |
| PP0_1 | 24,670,886 | 13,630,812 | 23 | 18,494,419 (22.7) | 6,039,689 (7.4) | 10,963,812 (3.1) | 99.4% | 99.2% | 99.2% |
| PP0_2 | 22,832,671 | 13,130,651 | 22 | 20,021,614 (24.6) | 6,554,371 (8.1) | 11,455,678 (3.4) | 99.4% | 99.2% | 99.1% |
| PP0_3 | 24,590,731 | 13,548,269 | 23 | 20,195,336 (24.4) | 6,682,634 (8.0) | 12,357,662 (3.6) | 99.4% | 99.2% | 99.1% |

The number of clean paired-end reads, mapped paired-end reads, the coverage, and the numbers of cytosines in all contexts are indicated. The percentages of methylated cytosines is indicated in parentheses. The conversion rates were calculated by aligning the reads to the lambda genome.

**Table S2.** Summary of RNA-seq reads statistics.

| **SampleID** | **Clean reads** | **Uniquely mapped reads** | **% Uniquely mapped reads** | **unmapped reads** | **% unmapped reads** |
| --- | --- | --- | --- | --- | --- |
| MM0_1 | 26,142,798 | 24,509,730 | 93.75% | 321,134 | 1.23% |
| MM0_2 | 26,275,497 | 24,753,733 | 94.21% | 291,783 | 1.11% |
| MM0_3 | 26,303,404 | 25,024,094 | 95.14% | 323,051 | 1.23% |
| MM6_1 | 25,811,650 | 24,631,637 | 95.43% | 393,538 | 1.52% |
| MM6_2 | 26,236,357 | 24,958,407 | 95.13% | 473,680 | 1.81% |
| MM6_3 | 26,302,916 | 25,201,992 | 95.81% | 417,488 | 1.59% |
| MM24_1 | 24,119,495 | 23,119,286 | 95.85% | 373,389 | 1.55% |
| MM24_2 | 26,298,388 | 25,108,774 | 95.48% | 434,838 | 1.65% |
| MM24_3 | 25,720,037 | 24,586,801 | 95.59% | 395,607 | 1.54% |
| MP0_1 | 26,301,855 | 24,333,831 | 92.52% | 424,369 | 1.61% |
| MP0_2 | 26,295,730 | 24,908,159 | 94.72% | 483,645 | 1.84% |
| MP0_3 | 26,286,139 | 24,866,739 | 94.60% | 349,476 | 1.33% |
| MP6_1 | 26,289,682 | 24,837,105 | 94.47% | 385,922 | 1.47% |
| MP6_2 | 26,147,476 | 24,873,762 | 95.13% | 460,211 | 1.76% |
| MP6_3 | 26,309,772 | 24,993,993 | 95.00% | 381,561 | 1.45% |
| MP24_1 | 26,298,616 | 24,899,328 | 94.68% | 438,273 | 1.67% |
| MP24_2 | 24,119,870 | 23,060,122 | 95.61% | 469,821 | 1.95% |
| MP24_3 | 25,631,943 | 24,366,457 | 95.06% | 552,908 | 2.16% |
| PM0_1 | 26,304,654 | 24,387,741 | 92.71% | 404,824 | 1.54% |
| PM0_2 | 26,178,695 | 24,728,890 | 94.46% | 375,670 | 1.44% |
| PM0_3 | 26,244,211 | 24,819,342 | 94.57% | 366,625 | 1.40% |
| PM6_1 | 26,265,498 | 24,884,454 | 94.74% | 417,270 | 1.59% |
| PM6_2 | 26,140,678 | 24,910,944 | 95.30% | 408,957 | 1.56% |
| PM6_3 | 26,306,406 | 25,193,882 | 95.77% | 396,569 | 1.51% |
| PM24_1 | 26,149,351 | 25,034,473 | 95.74% | 438,027 | 1.68% |
| PM24_2 | 26,305,382 | 25,256,896 | 96.01% | 417,147 | 1.59% |
| PM24_3 | 26,227,158 | 25,234,437 | 96.21% | 371,602 | 1.42% |
| PP0_1 | 26,290,203 | 24,519,620 | 93.27% | 335,879 | 1.28% |
| PP0_2 | 26,282,341 | 24,682,610 | 93.91% | 266,150 | 1.01% |
| PP0_3 | 26,278,123 | 24,632,828 | 93.74% | 533,676 | 2.03% |
| PP6_1 | 26,307,234 | 25,134,566 | 95.54% | 370,158 | 1.41% |
| PP6_2 | 26,297,542 | 25,095,152 | 95.43% | 385,644 | 1.47% |
| PP6_3 | 26,291,638 | 25,146,102 | 95.64% | 363,354 | 1.38% |
| PP24_1 | 26,139,931 | 24,990,889 | 95.60% | 444,423 | 1.70% |
| PP24_2 | 26,309,065 | 25,228,505 | 95.89% | 418,265 | 1.59% |
| PP24_3 | 26,286,857 | 25,189,547 | 95.83% | 444,390 | 1.69% |

**Table S3.** DMRs among the F1 descendants.

**Table S4.** Complete list of differentially expressed genes after treatment.

**Table S5.** Enriched GO terms of differentially expressed genes after treatment.

**Table S6.** Gene list of six clusters in Fig. S7B.

**Table S7.** Complete list of differentially expressed genes in different combinations.

**Table S8** Enriched GO terms of differentially expressed genes in different combinations.

